# The staphylococcal type VII secretion system delays inflammatory cell death during macrophage infection

**DOI:** 10.1101/2025.10.30.685513

**Authors:** Richard D. Allen, Robeena Farzand, Giridhar Chandrasekharan, Kate E. Watkins, Pooja Agarwal, Meera Unnikrishnan

## Abstract

The human pathogen *Staphylococcus aureus* is a facultatively intracellular bacterium that can replicate and survive within macrophages. While macrophage-mediated immune modulation is pivotal in *S. aureus* pathogenesis, bacterial manipulation of macrophage pathways remains poorly understood. The specialised type VII secretion system (T7SSb) is an important virulence-associated factor during staphylococcal infection, although, unlike its mycobacterial counterparts, its role in controlling staphylococcal-macrophage interactions remains unclear. Employing high-resolution time-lapse imaging, we demonstrated an induction of pore-induced cellular traps (PITs) during *S. aureus* infection of macrophages *in vitro*, with a higher number of PITs forming in macrophages infected with *S. aureus* lacking EssC (Δ*essC*), a central T7SS transporter. Interestingly, Δ*essC*-infected macrophages displayed increased bacterial escape compared to wild type (WT)-infected cells, implicating a role for T7SS in delaying macrophage cell death. Changes in cell death were accompanied by a reduction in efferocytosis of Δ*essC*-infected cells *in vitro*. Investigation of cell death signalling markers in WT and Δ*essC*-infected macrophages demonstrated that the T7SS delayed induction of key cell death pathways, necroptosis and pyroptosis, and a dual RNAseq analysis of WT- and Δ*essC*-infected macrophages indicated a T7SS-dependent modulation of bacterial oxidative stress and selected host chemokines. We also demonstrated that T7SS effectors, EsxA and EsxC individually delayed macrophage cell death. Interestingly, Δ*esxC* mutants demonstrated accelerated calcium fluxes, indicating that the T7SS-mediated suppression of cellular calcium signalling may cause the delay in cell death. Furthermore, in a murine skin infection model, we showed that effector and transporter mutants have distinct *S. aureus* infection outcomes. Our findings suggest for the first time a key role for T7SSb proteins in controlling macrophage cell death, which impacts staphylococcal survival within the host during infection.

**Highlights:**

- *S. aureus* T7SS delays formation of pore-induced traps in macrophages and bacterial escape
- T7SS dampens necroptosis and pyroptosis signalling in infected macrophages
- T7SS effectors EsxA and EsxC when expressed in macrophages delay cell death
- An effector EsxC shapes the timing of Ca^2+^ signalling in macrophages
- T7SS mutants have distinct outcomes in *S. aureus in vivo* infection

## Introduction

*Staphylococcus aureus* is a major healthcare and community-associated pathogen, responsible for millions of deaths annually around the globe^1^. This opportunistic pathogen and commensal can cause a wide array of infections from mild skin abscesses to serious pneumonia^2,3^. *S. aureus* has been demonstrated to invade, replicate and persist within a variety of host cells in recent years^4–6^. The ability to use the host cell as a survival niche is critical to its success as a pathogen and in evading antibiotics^7^. *S. aureus* employs several factors to control cell invasion, phagosomal disruption, intracellular replication and cytosolic persistence, in addition to the diverse arsenal of virulence factors to subvert a range of host immune factors^5,6,8^.

Macrophages are amongst the first cells that respond to *S. aureus* infections and are considered central for bacterial clearance through phagocytosis^3,9–11^. However, *S. aureus* survives, replicates, and escapes from macrophages into the extracellular environment, facilitating the propagation of infection^3,12,13^. The intracellular environment of macrophages provides a desirable niche for *S. aureus* persistence, as evidenced by its ability to persist for over 24 hours in alveolar macrophages^9,14^. Macrophages also serve as a ‘Trojan horse’, aiding in the dissemination of *S. aureus* to distal sites, seeding lesions in vital organs^12,15^. Thus, macrophage dynamics and clearance can significantly influence infection outcomes^5,16,17^. Nevertheless, intracellular infection of macrophages often triggers programmed cell death to clear the pathogen^18^.

*S. aureus* exhibits remarkable proficiency in the manipulation of major cellular death programmes, apoptosis, necroptosis and pyroptosis, across diverse host cells including macrophages^19–22^. The bacterium has been reported to both promote and inhibit inflammatory cell death, pyroptosis, exploiting this pathway for intracellular persistence^23–26^. A few staphylococcal factors such as α-toxin, leukocidins, and phenol-soluble modulins have been reported to induce host cell death^20,27–29^, however, little is known about how *S. aureus* delays cell death. We and others have reported previously that EsxA, an effector of the type VII secretion system (T7SS), was shown to delay apoptosis during intracellular epithelial and dendritic cell infection, promoting host cell survival^30,31^.

The staphylococcal T7SS contains a membrane-associated FtsK/SpoIIIE ATPase protein, EssC, which is the central transporter that exports several effector proteins including WXG100 motif containing proteins EsxA and EsxB, small proteins EsxC, EsxD, and toxins like EsaD, TspA and TelA^32–36^. The staphylococcal T7SS is heterogeneous and modular in nature with the module 1 conserved in all commonly studied strains, including USA300^37^. Whilst the exact functions of the system during infection remain elusive, recent discoveries include its involvement in inter-species competition via a toxin-antitoxin pair, as well as potential roles in iron acquisition and bacterial membrane homeostasis^34,38–41^. Additionally, the T7SS is required for staphylococcal virulence and persistence in several models of infection^32,34,36,42^ Click or tap here to enter text.. The system further appears to be immunostimulatory with effectors EsaE and EsaD stimulating IL-1β and IL-12 cytokine production during infection^43,44^. Consistent with these observations, Type VII proteins have been implicated in human disease, with T7SS genes upregulated in prosthetic joint infections and in human cystic fibrosis patients^45,46^. Nevertheless, the molecular targets of the staphylococcal T7SS effectors in the host remain unknown.

Here, we investigate the role of T7SS in *S. aureus*-macrophage interactions. Employing an *S. aureus* mutant lacking EssC, a central T7SS transporter, and high-resolution live imaging we demonstrate that the T7SS delays the formation of *S. aureus-*induced pore induced traps (PITs) in macrophages, impacts bacterial release from macrophages and affects efferocytosis of infected cells. We show that the T7SS delays the induction of inflammatory cell death pathways, pyroptosis and necroptosis in macrophages during infection. A dual RNA-seq further revealed that local chemokine responses were subsequently modulated by the T7SS.

Specific T7SS effectors modulated levels of macrophage cell death, and effector-specific effects were evident during infection *in vivo*. We demonstrate that a T7SS effector alters the timing of calcium signalling in infected macrophages, which may contribute to delayed cell death. Thus, our data indicate a central role for the *S. aureus* T7SS proteins in delaying *S. aureus* induced macrophage cell death.

## Results

### *S. aureus* induces the formation of pore-induced trap-like structures during macrophage infection, with an increase in PIT formation in the absence of T7SS

To investigate the role of the T7SS during macrophage infection, we first examined *S. aureus*-infection in macrophages using high-resolution time-lapse microscopy. During intracellular *S. aureus* USA300 JE2 infection of THP-1 macrophages a heterogeneous population of cellular morphologies was observed, including ‘spindle’, spread out ‘fried egg’, and smaller ‘rounded’ (Figure 1A) shapes with a small subset of infected macrophages undergoing apoptotic-like death characterised by cellular blebbing (Figure 1A). Interestingly, we observed the emergence of pore-induced intracellular trap (PIT)-like structures^47^ with the cell ballooning into a spherical structure, and with the other morphologies (spindle, fried egg and rounded) declining over time (Figure 1A).

**Figure 1.**
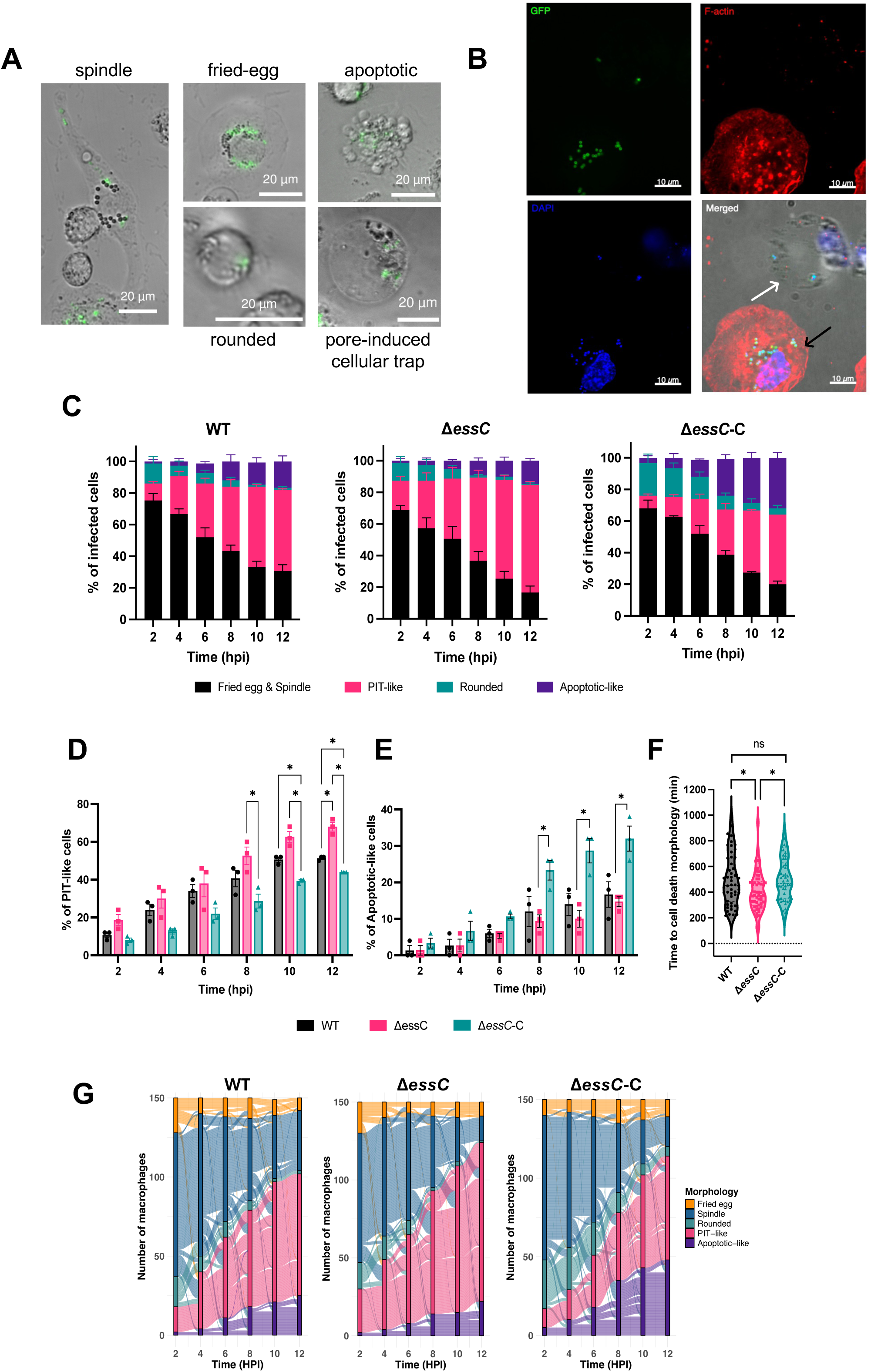
*S. aureus* induction of PIT-like structures during macrophage infection with a T7SS-associated decrease in PIT formation. THP-1 macrophages and bone marrow derived macrophages (BMDM) infected with YFP-expressing *S. aureus* JE2 WT at MOI 10:1 followed by extracellular killing of bacteria using lysostaphin and gentamicin. A heterogeneous population of BMDMs was observed during *S. aureus* infection including elongated spindle, spread out fried egg, rounded, apoptotic-like and pore-induced cellular trap (PIT)-like structures, scale bar represents 20 μm **(A).** Data are representative of N > 3. **B**) WT-infected THP-1 macrophages at 4 hpi stained for F-actin (red), DAPI (blue), and vancomycin-BODIPY (green), black arrow indicates intact macrophage, and white arrow indicates PIT structure, scale bar represents 10 μm (N = 2). BMDMs infected with YFP-expressing *S. aur*eus JE2 WT and Δ*essC* for 1 hr at MOI 10:1 before extracellular killing of bacteria. Individually infected macrophages were quantified blinded from time-lapse microscopy. **C**) The percentage of infected macrophage morphologies over time. N = 3, mean ± standard deviation (SD) Quantification of PIT-like structures (**D**) and apoptotic-like (**E)** morphology over time in infected BMDMs. N = 3, mean ± SD. **F**) Time to cell death morphology (apoptotic and PIT-like) of infected BMDMs, N = 3, median with quartiles presented. **G)** The morphological fate of individually infected macrophages over 12 hpi for WT, Δ*essC*, and Δ*essC-C* infection. Statistical significance was calculated for D-F by two-way ANOVA and Tukey’s multiple comparison test *P<0.05, **P<0.01, ns = non-significant.

We observed that the PIT-like structures induced during infection contained *S. aureus*. Bacteria could replicate, lyse and form microcolonies within these structures (Figure S1A), but mostly, they remained within these structures without replication over time (Figure S1B), suggesting that the PIT-like structures may spatiotemporally entrap intracellular *S. aureus*. Confocal microscopy to study cellular changes associated with PIT formation revealed contrasting cellular morphology between infected but viable macrophages and the PIT-like structures, both containing intracellular *S. aureus* (Figure 1B). The actin cytoskeleton was completely absent in these PIT-like structures, with the nucleus condensed to one section of the corpse. Upon *S. aureus* infection, the PIT-like structures displayed condensed nuclei and a reduction in soluble lipid granules in contrast to lipid granules dispersed throughout the cytoplasm in uninfected THP-1 macrophages (Figure S1C). Consistent with previous findings^48^, our observations indicate these structures exhibit permeabilised plasma membranes without complete rupture, resulting in a loss of soluble contents and retention of insoluble organelles like the nucleus^47^.

Next, we carefully quantified the morphologies observed during infection (Fig 1A). *In vitro* infection profiles of a wild type (WT) *S. aureus* JE2 strain was compared to a mutant lacking the central T7SS transporter EssC during infection of bone marrow derived macrophages (BMDMs). Morphological changes of 50 infected cells in each field of view (blinded samples, N = 3), were manually tracked at various time points. PITs become the dominant morphology by the end of the infection with a small number of apoptotic-like alongside fried egg and spindle morphologies (Figure 1C). Uninfected macrophages largely displayed ‘spindle’ or ‘fried egg’ morphologies with a small proportion of cells showing ‘rounded’ morphologies (Figure S1D). Interestingly, by 12 hours post infection (hpi) Δ*essC*-infected BMDMs displayed a significant increase in the formation of PITs, whilst a complemented strain, where *essC* was episomally expressed in Δ*essC* (Δ*essC*-C), exhibited a decrease in these (Figure 1D). Δ*essC*-C infected BMDMs instead displayed increased apoptotic-like morphologies by 12 hpi compared to Δ*essC*-infected BMDMs (Figure 1E). Assessment of overall time taken to visualise morphologies associated with cell death revealed a peak of cell death ∼500 minutes post infection for WT (Figure 1F), which was significantly accelerated during Δ*essC*-infection to ∼400 minutes and reversed in Δ*essC*-C-infected macrophages. Macrophage cytotoxicity was also assessed by a lactate dehydrogenase release assay, but this was unable to detect differences between WT- and Δ*essC*-infection in THP-1 macrophages (Figure S2).

Tracking the morphological fates of individual infected macrophages revealed that during WT-infection, PITs were predominately formed from cells with spindle morphologies rather than cells with rounded and fried egg morphologies (Figure 1G). All cells displaying ‘alive’ morphologies formed apoptotic-like morphologies later, with spindle morphologies being the main contributor. Cells infected with Δ*essC* and Δ*essC*-C mainly displayed differences in the proportions of PIT-like and apoptotic-like morphologies compared to WT-infected cells, although there were little differences to the overall dynamics of morphological fate over time.

### T7SS impacts bacterial escape from *S. aureus*-infected macrophages

During intracellular macrophage infection, intracellular *S. aureus* escapes into the extracellular environment, often forming microcolonies on the cell layer^31,48^. Notably, at 12 hpi Δ*essC*-infected macrophages exhibited a greater number of microcolonies compared to WT-infected during both BMDM and THP-1 macrophage infection (Figure 2A, Figure S3), suggesting the T7SS influences bacterial escape. The time to microcolony formation was assessed by measuring time to cell lysis and subsequent bacterial release from cells. Correspondingly, Δ*essC*-infected macrophages displayed an accelerated time to microcolony formation and greater number of individual macrophages which lysed to establish microcolonies (Figure 2B, C). As expected, the phenotypes were reversed with Δ*essC*-C. These results suggest that the T7SS contributes to *S. aureus* escape from infected macrophages.

**Figure 2.**
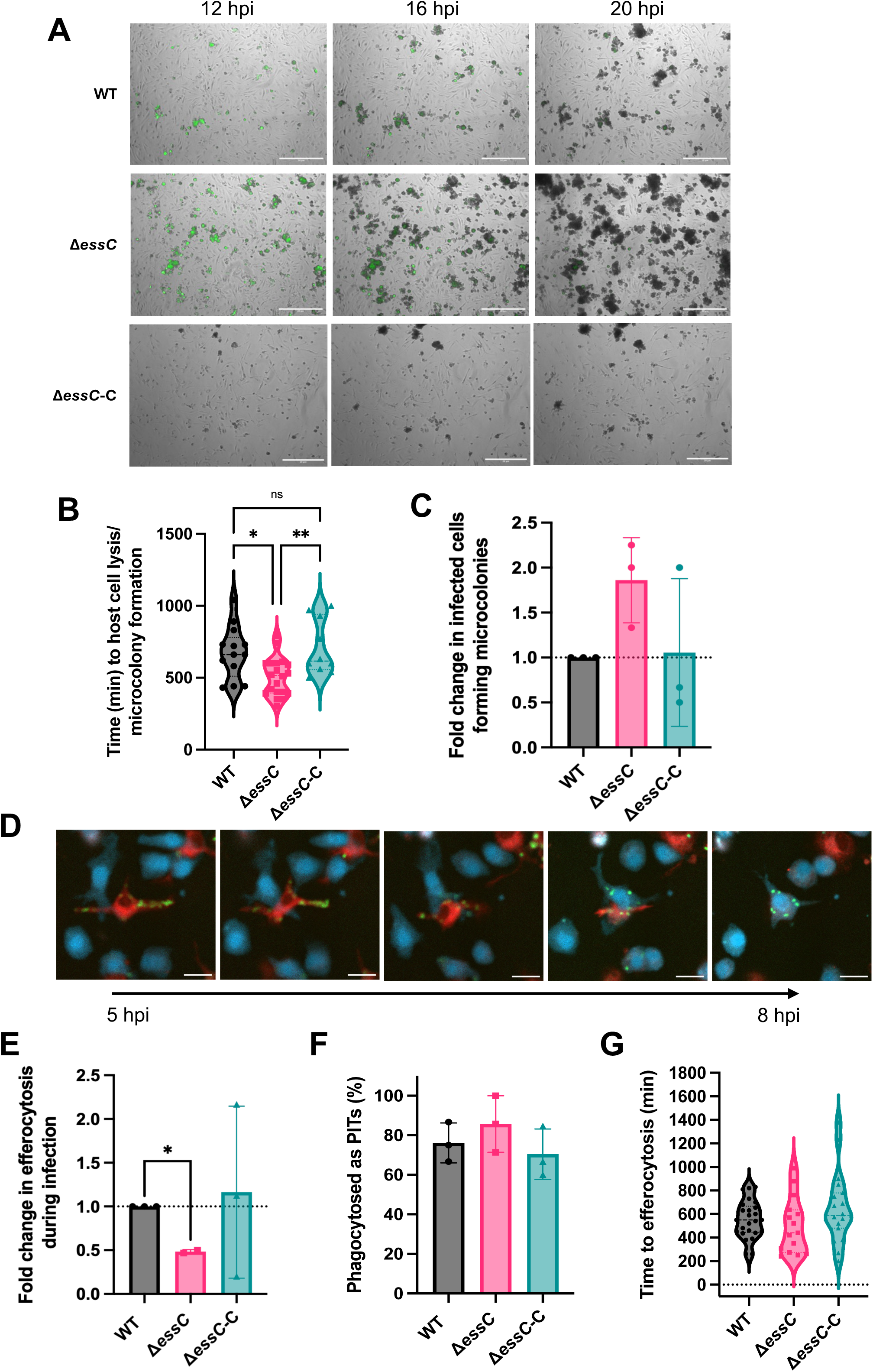
T7SS impacts *S. aureus* escape from macrophages and promotes efferocytosis of infected macrophages. BMDMs infected with YFP-expressing *S. aureus* JE2 WT and Δ*essC* for 1 hr at MOI 10:1 before extracellular killing of bacteria. **A)** Representative images of *S. aureus* escaping BMDMs to form microcolonies on the cell layer from 12-20 hpi, scale bar represents 20 μm. **B)** Time to infected host cell lysis and subsequent microcolony formation (min), N = 3, median presented with quartiles. *P<0.05, **P<0.01, ns = non-significant by one-way ANOVA with Tukey’s multiple comparison test. **C**) Infected macrophages counted to lyse and form subsequent microcolonies, presented as relative fold change to WT. N = 3, mean ± SD, P value ns by one sample t-test. **D**) iBMDMs were stained with CellTracker^TM^ CMPTX or CMAC prior to infection as described in Methods. Representative images of efferocytosis of YFP-expressing JE2 WT (green), infected iBMDM (red) and uninfected iBMDM (blue) across 5-8 hpi during infection, imaged by live confocal microscopy, scale bar represents 5 μm. **E**) BMDMs were infected with YFP-expressing JE2 WT, Δ*essC*, and Δ*essC-C* strains and imaged by time-lapse microscopy, quantification was carried out blinded by counting 50 infected cells per field across replicates: the number of efferocytosis events for infected BMDMs presented as relative fold change to WT-infected, one-sample t-test against WT, N = 3, mean ± SD *P < 0.05. **F**) Quantification of the percentage of infected macrophages efferocytosed as PIT-like structures, N = 3, mean ± SD, ns by two-way ANOVA. **G**) Quantification of time to efferocytosis events, median presented with quartiles, N = 3, ns by two-way ANOVA with Tukey’s multiple comparisons test.

### T7SS promotes efferocytosis of infected macrophages

Infected and dying cells are usually removed by phagocytosis termed efferocytosis. To study if *S. aureus*-infected macrophages were efferocytosed by uninfected macrophages, we visualised efferocytosis by live confocal time-lapse microscopy using macrophages labelled with different CellTracker dyes as described in Methods. An efferocytosis event was defined as when the uninfected macrophage (CMAC stained) had engulfed the infected macrophage (CMPTX stained). Efferocytosis events for infected macrophages both with ‘alive’ morphologies and PIT corpses were observed (Figure 2D). Quantification of these events revealed that Δ*essC*-infected BMDMs were efferocytosed significantly less than WT-infected cells (Figure 2E). Interestingly, >75% of efferocytosis events across strains were of PITs suggesting that they may be important to infection dynamics and clearance (Figure 2F). The T7SS did not significantly impact the timing of efferocytosis events which averaged ∼500-600 min post infection (Figure 2G), although a slight but statistically non-significant increase in efferocytosis events of PITs was observed (Figure 2F). Thus, the *S. aureus* T7SS may be involved in promoting efferocytosis of infected macrophages.

### T7SS modulates necroptosis in infected macrophages

Inflammatory cell death pathways have been reported to be activated by *S. aureus-*infected macrophages^49,50^. Additionally, PITs were shown to be associated with inflammatory cell death^47,51^. As we observed low levels of apoptotic-like cell death morphologies in our infection assays and as necroptosis is generally a back-up mechanism to apoptosis inhibition^52^, we hypothesised that the necroptosis pathway was impacted by T7SS proteins during macrophage infection. We observed a minor decrease in CFU in WT- and Δ*essC*-infected BMDMs (Figure S4A, B), and a significant decrease in WT PIT formation until 10 hpi (Figure S4C) upon treatment of cells with the necroptosis inhibitor necrosulfonamide (NSA), which blocks recruitment and phosphorylation of the mixed lineage kinase domain-like (MLKL) executioner by the necrosome complex^53^. Morphological differences induced by NSA treatment were observed independent of the T7SS in THP-1 macrophages (Figure S5 A-F). Remarkably, the initial wave of cell death morphology and corresponding cytotoxicity was significantly delayed upon NSA treatment during WT-infection (Figure 3A, B), but not in Δ*essC*-infected macrophages.

**Figure 3.**
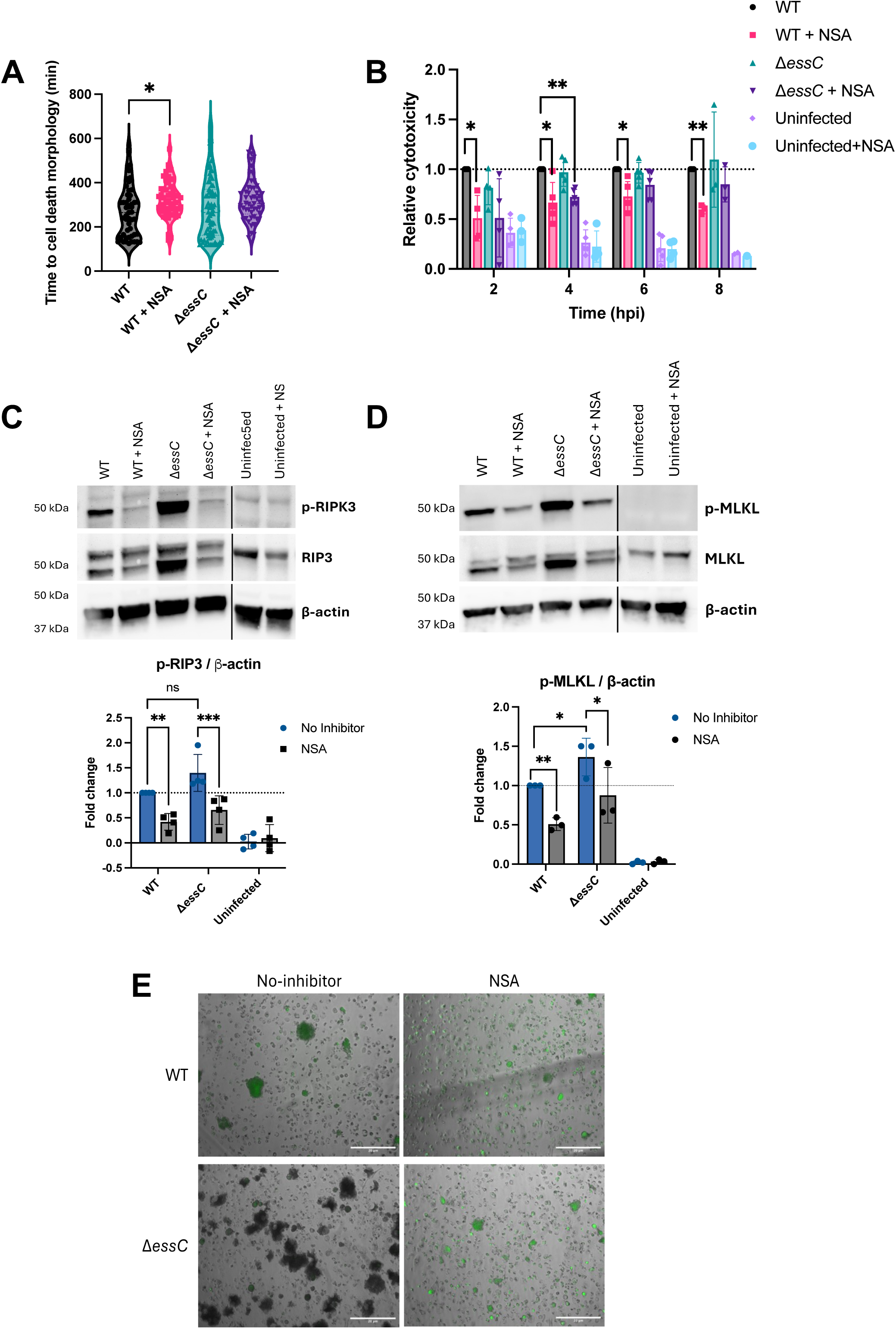
Necroptosis drives initial cell death of macrophages during *S. aureus* infection. **A**) Violin plots display time to cell death morphology (PIT-like and apoptotic-like) for JE2 WT- and Δ*essC*-infected THP-1 macrophages with or without necrosulfonamide (NSA) treatment, N = 3, median presented with quartiles, *P < 0.05 and **P < 0.01 by Kruskal-Wallis test with Dunn’s multiple comparisons test. **B**) The relative cytotoxicity to the WT measured by LDH release during THP-1 macrophage infection, N = 3, mean ± SD, one-sample t-test relative to WT. *P < 0.05 and **P < 0.01. THP-1 macrophage lysates obtained at 4 hpi were probed with anti-RIP3, anti-phosphorylated-RIPK3 (p-RIPK3) (**C**) anti-MLKL, phosphorylated-MLKL (p-MLKL) (**D**) and with loading control of anti-β-actin. Representative immunoblots are shown. The relative quantification of bands by normalisation to loading control, presented as relative fold change to WT no-inhibitor infection is shown below each image. N = 3, mean ± SD, *P < 0.05 and **P < 0.01 by one-sample t-test. (**E**) Representative images of microcolonies on infected THP-1 macrophages at 10 hpi, with or without NSA treatment, scale bar represents 20 μm, N = 3.

Immunoblotting of key necrosome components, receptor interacting protein kinase 3 (RIPK3) and the executioner MLKL at 4 hpi, revealed increased phosphorylation of RIPK3 and MLKL in Δ*essC*-infected macrophages compared to WT-infected macrophages (Figure 3C, D). These data suggest that the T7SS attenuates necroptosis induction during macrophage infection. As expected, NSA treatment reduced p-RIPK3 and p-MLKL levels during WT and Δ*essC*-infection.

Intriguingly, NSA treatment led to a reduction in the formation of microcolonies during infection with either strain, implicating necroptosis in extracellular escape (Figure 3E). LIVE/DEAD bacterial staining revealed that the microcolonies predominantly were comprised of damaged bacteria as indicated by red PI staining (Figure S6A). Although fewer microcolonies were seen after NSA treatment, they contained more *S. aureus* that were intact (green SYTO-9 staining). Furthermore, intracellular *S. aureus* retrieved from macrophages by cell lysis, when stained with LIVE/DEAD stain were heterogeneous with live and damaged bacteria (Figure S6B), and correlated with a significant reduction in colony counts upon re-infection of fresh macrophages (Figure S6C). However, no differences were observed between WT- and Δ*essC* re-infections. Thus *S. aureus* appears to experience considerable stress within macrophages, induced by necroptosis, which affects downstream escape and microcolony formation. Overall, the data suggest that *S. aureus*-induced necroptosis is delayed by T7SS during macrophage infection.

### T7SS causes a delay of pyroptosis in *S. aureus-*infected macrophages

As formation of PITs was still observed when necroptosis signalling was inhibited, we tested the involvement of the other major inflammatory death pathway, pyroptosis, which is known to contribute to cell death during infection^20,47,54^. Immunoblotting for pyroptosis executioner gasdermin D (GSDMD) revealed at 4 hpi a trend of increased accumulation and significantly greater cleavage (c-GSDMD) in Δ*essC*-infected macrophages when compared to WT-infection (Figure 4A). The observed increase in GSDMD signalling is largely restored to WT levels by 6 hpi.

**Figure 4.**
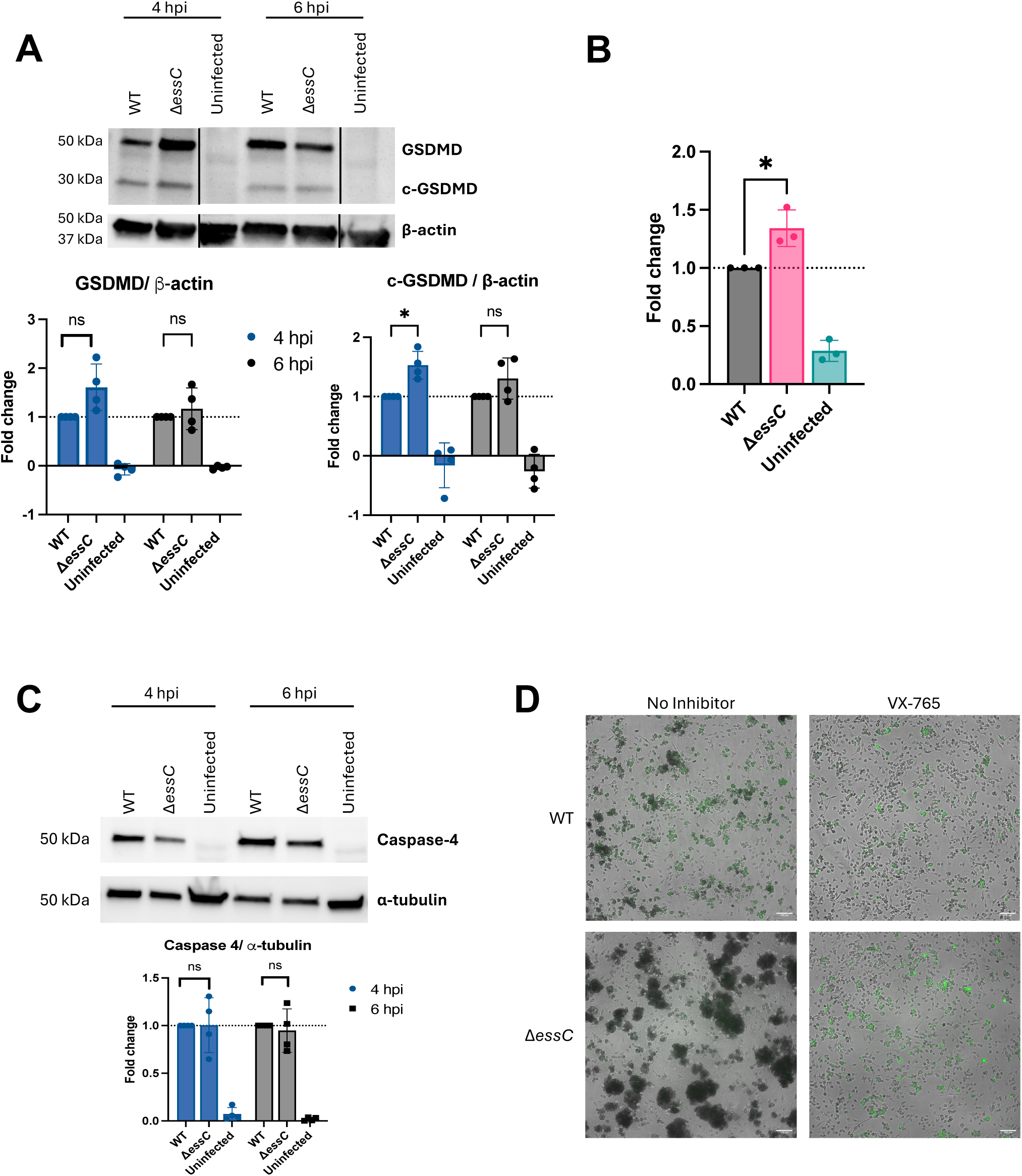
Staphylococcal T7SS modulates canonical pyroptosis during macrophage infection. THP-1 macrophages and BMDMs were infected with *S. aureus* JE2 WT and Δ*essC* for 1 hr at MOI 10:1 before extracellular killing of bacteria. **A**) THP-1 macrophage lysates obtained at 4 and 6 hpi were probed with anti-gasdermin (GSDMD), anti-cleaved-GSDMD (c-GSDMD), and loading control anti-β-actin. A representative immunoblot is shown. Quantification of bands on immunoblots by normalisation to loading control is shown below the image, presented as relative fold change to WT at each time-point for GSDMD and c-GSDMD. N = 3, mean ± SD, one-sample t-test relative to WT, *P < 0.05 and **P < 0.01. **B**) Caspase-1 activation as measured by Caspase-1 Glo luminescence at 2 hpi presented as relative light units (RLU) and fold change relative to WT, N = 3, mean ± SD, one-sample t-test relative to WT, *P < 0.05. **C**) Representative immunoblots of Caspase-4 with loading control α-tubulin from THP-1 macrophage lysates obtained at 4 and 6 hpi. Relative quantification of Caspase-4 bands normalised to loading control is shown below the image, presented as fold change relative to WT. N = 3, mean ± SD, differences were ns by one-sample t-test. **D**) Representative images of microcolony formation at 16 hpi for YFP-expressing strains of JE2 WT- and Δ*essC*-infected iBMDMs with and without caspase inhibitor VX-765, scale bar represents 100 μm, N = 2.

Upstream of the pyroptosis executioner, Caspase-1 activation was greater in Δ*essC*-infected macrophages at 2 hpi (Figure 4B), suggesting the T7SS may act at the inflammasome level. However, no T7SS-mediated effects were seen on NLRP3, an inflammasome known to be activated by *S. aureus* virulence factors (Figure S7)^55,56^. Previous reports have highlighted the activation of the non-canonical murine Caspase-11 by *S. aureus*, mediated by reactive oxygen species (ROS)^57^. We observed the activation of Caspase-4, the human equivalent of Caspase-11, from 4 hpi in *S. aureus* infected macrophages, although this was also independent of the T7SS (Figure 4C).

As with NSA, introduction of the caspase inhibitor VX-765, reduced bacterial escape and subsequent microcolony formation during infection across both WT and Δ*essC* strains (Figure 4D). Thus, the T7SS dampens pyroptosis signalling, alongside similar effects to necroptosis signalling, suggesting that pyroptosis also contributes to individual macrophage fate during *S. aureus* infection.

### T7SS is associated with oxidative stress during macrophage infection

Cellular death signalling pathways interact closely with many other cellular signalling pathways and functions^58,59^. Hence, to gain further insight into T7SS-mediated effects, we determined T7SS-specific global transcriptional responses induced in the infected macrophages by dual RNA-seq. Bacterial and mammalian cell responses were compared between *S. aureus* WT- and Δ*essC*-infected THP-1 macrophages at 2 and 6 hpi (Fig S8).

Direct comparison of Δ*essC*-to WT-infected macrophages revealed 8 differentially expressed genes (DEGs) at 2 hpi and 20 DEGs at 6 hpi (Table S3). It was interesting to note that at 2 hpi, the enzyme inositol-tetrakisphosphate 1-kinase (ITPK1) involved in the synthesis of inositol phosphate which controls MLKL-mediated necroptosis^60^, was differentially upregulated in the Δ*essC*-infected macrophages. Pathway enrichment by the KEGG database revealed Δ*essC*-infected macrophages displayed up-regulation of inositol phosphate metabolism, peroxisome, and phosphatidylinositol signalling system and down-regulation of nucleotide sugar metabolism (Figure S9A-C). Further analysis identified unique DEGs within each condition, i.e. specific DEGs observed only in either WT-or Δ*essC*-infected macrophages when compared to uninfected macrophages. Unique DEGs of WT-infected macrophages displayed pathway enrichment for the PPAR signalling pathway which has been associated with a multifaceted role during macrophage inflammation and the response to microbes^61^ (Figure S9 D-F, Table S4). The chemokines *CCL4* and *CCL24* were significantly upregulated in WT-infected macrophages by 6 hpi whereas *CCL3* was significantly up-regulated in Δ*essC*-infected macrophages (Figure 5A). The cytokine *OSM* was significantly up-regulated in Δ*essC*-infected macrophages 2 hpi, although there were no significant changes between other cytokines, compared with uninfected macrophages (Figure S10A).

**Figure 5.**
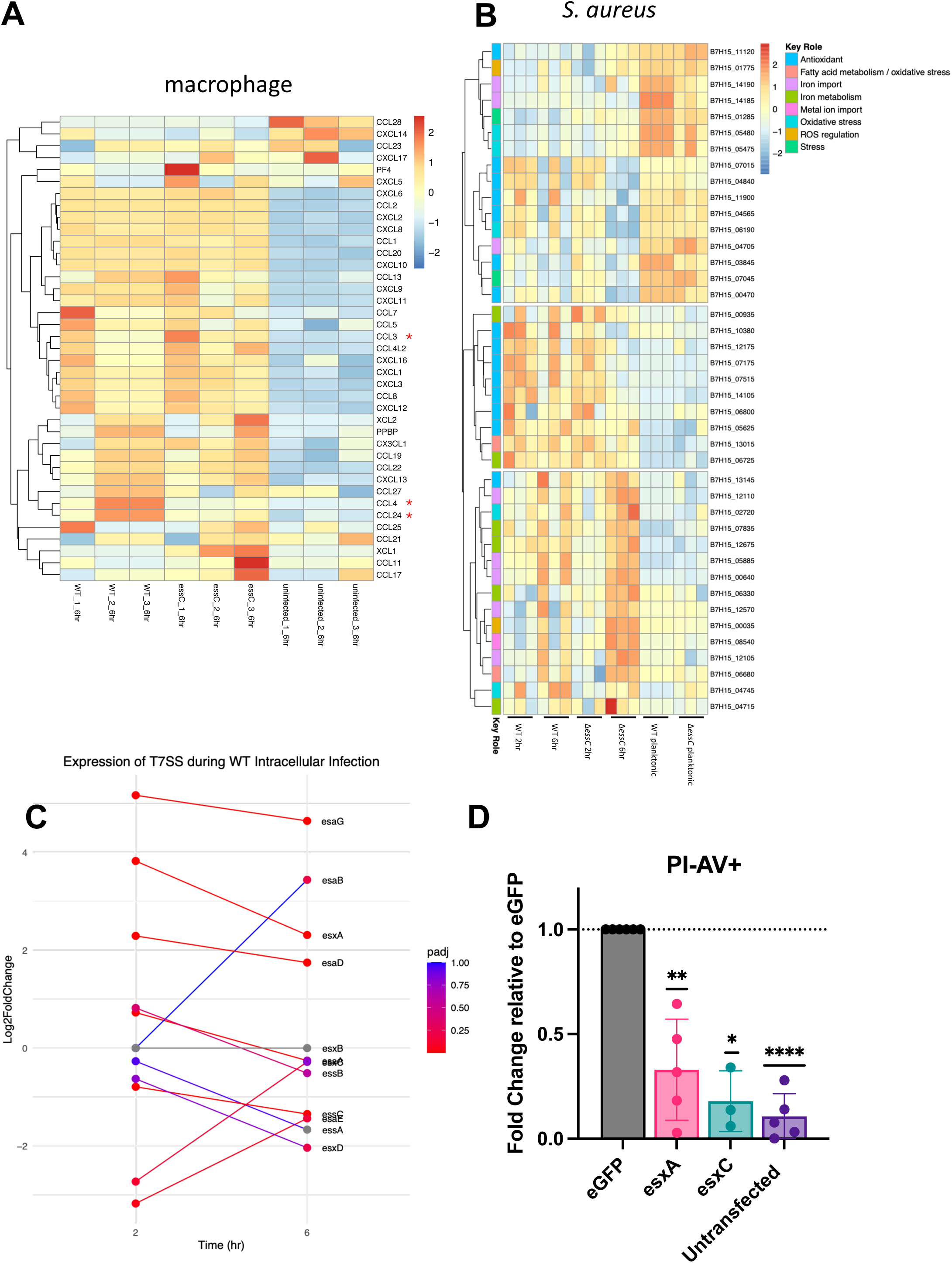
Staphylococcal T7SS genes are induced intracellularly, contribute to oxidative stress and delay macrophage cell death during intracellular macrophage infection. **A)** Heatmap of chemokine expression in JE2 WT or Δ*essC* infected THP-1 macrophages and uninfected macrophages, at 6 hpi. **B)** Normalised *S. aureus* counts from dual RNA-seq of *S. aureus* JE2 infected THP-1 macrophages highlighting differences in iron metabolism, oxidative stress, and ROS pathways in JE2 *S. aureus*. **C)** Relative expression of T7SS locus in WT JE2 *S. aureus* during intracellular macrophage infection compared against planktonic JE2 WT. **D) i**BMDMs were transfected with either pEGFP-N1, pEGFP-N1+*esxA*, or pEGFP-N1+*esxC* via nucleofection. At 12 hrs post transfection, macrophages were stained with Annexin V (AV) Blue Pacific and propidium iodide (PI) and analysed by flow cytometry. Transfected cells were gated for eGFP expression. The number of PI- and AV+ macrophages relative to the empty control vector (eGFP). N = 3, mean ± SD and one-sample t-test, *P < 0.05, **P < 0.01, ***P < 0.001, and ****P < 0.0001.

Direct comparison of intracellular *S. aureus* Δ*essC* to WT revealed 4 DEGs at 2 hpi and 6 DEGs at 6 hpi (Table S5). The cold-shock proteins CspC and CspB and the virulence factor clumping factor B were upregulated in intracellular Δ*essC.* Non-biased hierarchical clustering of the microbial DEGs also revealed 3 clusters displaying distinct expression profiles in intracellular WT and Δ*essC*, compared to planktonic *S. aureus*. Intracellular macrophage WT-infection induces an up-regulation of iron metabolism, as well as antioxidant and oxidative stress pathways, suggesting the pathogen is adapting to the nutrient-depleted environment within the macrophage (Figure 5B; Table S4, Table S6). By 6 hpi, this profile is inversed, wherein intracellular Δ*essC* displayed both up- and down-regulation of genes associated with these pathways, suggesting that the T7SS mutant was delayed in responding to the intracellular macrophage environment.

Several DEGs involved in the ROS and iron-dependent cell death pathway ferroptosis, were differentially regulated between WT- and Δ*essC*-infected macrophages (Figure S10B). An inducer of ferroptosis, *ACSL4*, was up-regulated during both WT- and Δ*essC*-infection at 6 hpi, however, only Δ*essC*-infected macrophages exhibited significant up-regulation of *ECH1*, an inhibitor of ferroptosis at 2 hpi^62^ (Figure S10B; Table S7). WT-infected macrophages also showed a down-regulation of *SLC3A2*, essential for cysteine import and inhibiting ferroptosis^63^. However, there was no significant difference in levels of antagonist regulator glutathione peroxidase (GPX4)^64^ between WT-infected and Δ*essC*-infected macrophages, indicating a T7SS-independent modulation of ferroptosis (Figure S10C), and suggesting involvement of other staphylococcal factors in *S. aureus*-induced ferroptosis during infection.

### T7SS effectors impact cell death in macrophages

As the T7SS encodes numerous effectors that are exported, presumably also into the host cell, we sought to elucidate the impact of individual T7SS effectors upon macrophage cell death. We observed in the RNA-seq analysis a significant increase in expression of the T7SS secreted effectors EsxA and EsaD (nuclease) and the anti-toxin EsaG, during intracellular *S. aureus* infection in macrophages compared to expression in planktonic growth, although no change in expression was observed for the transporter EssC itself (Figure 5C, Figure S11A). To test the direct activity of effectors within cells, we transfected staphylococcal *esxA* and *esxC* (expressed upstream of eGFP on a peGFP-N1 plasmid under a CMV promoter) individually into immortalised BMDMs (iBMDMs). At 12 hours post-transfection when GFP expression was strongest, cells were stained for annexin V (AV, marker for apoptosis) and propidium iodide (PI, marker for necrosis) and analysed by flow cytometry. Both EsxA and EsxC significantly reduced the number of apoptotic cells (PI-, AV+) in the population compared to the no effector control (Figure 5D, Figure S11B). Although cells transfected with EsxC appeared to increase numbers of PI+AV+- and PI+AV-populations, there was no significant difference compared to the control across multiple experiments (Figure S11C, D). Therefore, our data suggest that individual T7SS effectors can directly suppress intracellular cell death signalling in macrophages.

### T7SS effector EsxC alters Ca^2+^ signalling in infected macrophages

Calcium is known to be a key regulator of cell death, with clear links to pyroptosis and necroptosis^65,66^. As individual T7SS effectors directly influence macrophage cell death, we next examined whether they modulate intracellular Ca²⁺ signalling dynamics during infection. Ca²⁺ flux during infection was first assessed using a fluorescent calcium probe Fluo-8, to quantify Ca²⁺ at a population-level, within the entire field of view. WT infection produced a steady rise in cumulative Ca²⁺ signal over 120 min (Figure 6A). Δ*essC* infected cells showed reduced Ca²⁺ fluorescence intensities, with the complemented strain Δ*essC-C* restoring cumulative Ca²⁺ responses to WT levels. The *S. aureus* mutant lacking the effector EsxC, Δ*esxC*, displayed a markedly reduced fluorescence intensity compared to the WT, whilst a USA300 strain lacking EsxA, Δ*esxA,* did not show a significant difference (Figure S12). A reduced overall calcium signal in mutant-infected cells may be due to loss of cellular Fluo-8 from perforated cells due to earlier cell death.

**Figure 6.**
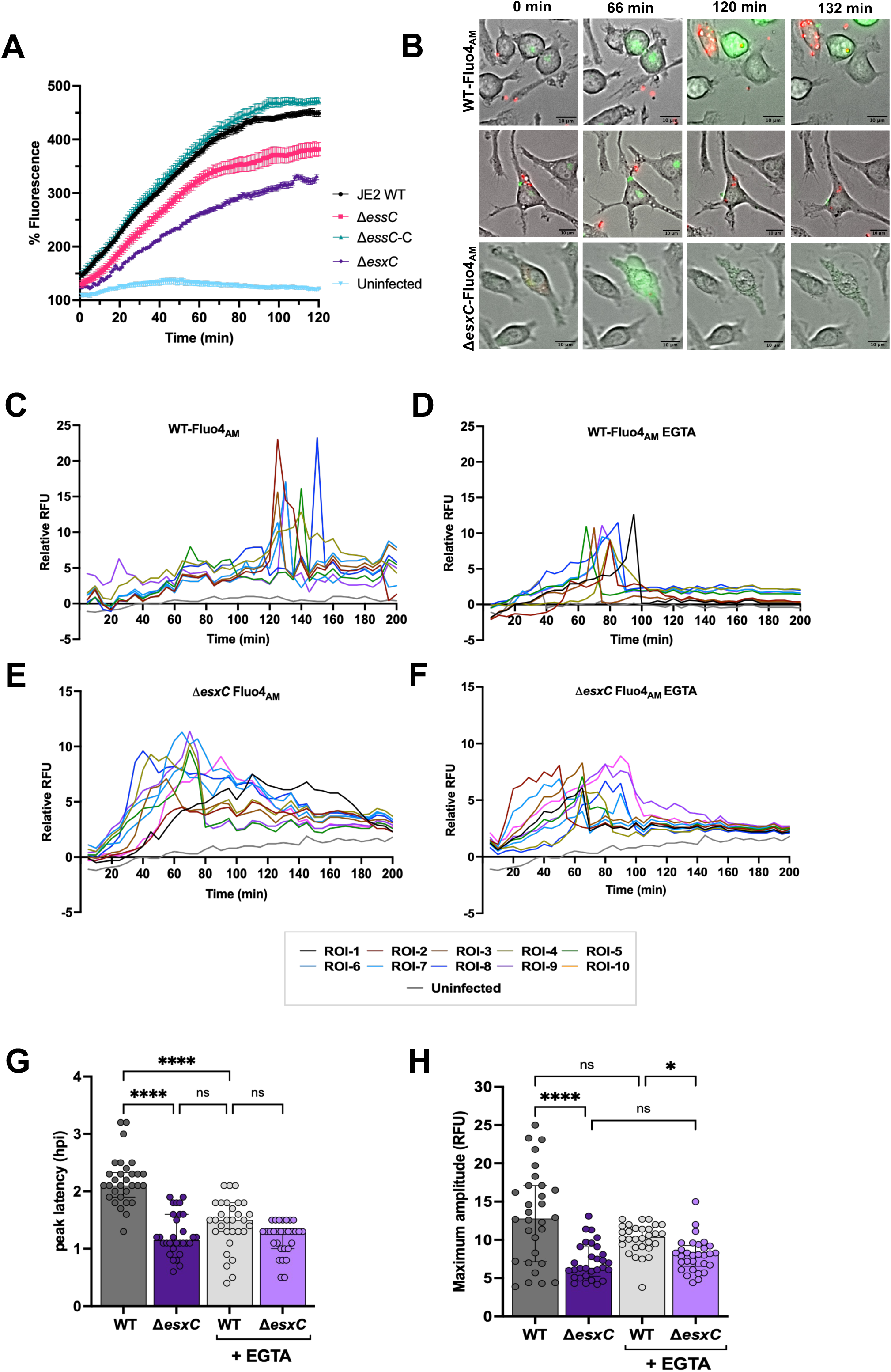
T7SS effector EsxC modulates intracellular Ca²⁺ signalling dynamics in macrophages during infection. (**A**) For measuring intracellular calcium levels following infection, macrophages were loaded with Fluo-8-AM and infected with WT (JE2), Δ*essC*, Δ*essC-C* and Δ*esxC* strains at MOI 10:1 or left uninfected. Time-lapse fluorescence imaging was performed at 1 min intervals for 120 min. For each time point, fluorescence intensity was quantified across the entire field of view, cumulative % fluorescence = (ΔF/F₀) × 100 relative to baseline, N = 3, mean ± SEM. For single cell Ca^2+^ signalling, iBMDMs were loaded with Fluo-4-AM and infected with WT (JE2) or Δ*esxC* mutant at MOI 10:1. (**B**) Representative time-lapse images of Fluo-4-AM fluorescence in infected iBMDMs illustrating distinct intracellular Ca²⁺ signalling dynamics during infection, green fluorescence denotes intracellular Ca²⁺, scale bars = 10 μm. Intracellular Ca²⁺ flux was monitored over time in single WT- or Δ*esxC*-infected, Fluo-4-AM treated macrophages (**C**, **E**). Following Fluo-4-AM loading, cells were treated with 3 mM EGTA to chelate extracellular Ca²⁺ prior to infection with WT or Δ*esxC* (**D**, **F**). For **C-F**, fluorescence intensities were quantified for individual ROIs corresponding to single infected cells, background-subtracted, and normalised to the initial 0–20 min baseline to obtain relative fluorescence values, data shown are representative of one of 3 biological replicates. (**G**) Peak latency is the time point of maximum amplitude (hpi), measured from 30 single infected cells from 3 independent experiments, median ± interquartile range (IQR) shown. (**H**) The maximum amplitude was determined as the highest relative fluorescence value of 30 single infected cells from 3 independent experiments, median ± IQR shown. Statistical significance (**G**, **H**) was determined using a Kruskal-Wallis test with Dunn’s multiple comparisons test, *P < 0.05, ****P < 0.0001.

To investigate this further at a single-cell level, intracellular Ca²⁺ flux was assessed using the fluorescent calcium probe Fluo-4, by measuring the amplitude of Ca²⁺ flux in individual infected macrophages. During WT infection, macrophages showed a characteristic Ca²⁺ peak at median 2.1 (IQR 1.9 - 2.32) hpi, indicating that WT *S. aureus* triggers a gradual build-up of Ca²⁺ mobilisation (Figure 6B, C, G, H). When extracellular Ca²⁺ was chelated using EGTA, this timing shifted and Ca²⁺ peaks occurred earlier (Figure 6 B, D, G, H) compared to untreated WT-infected cells, indicating that extracellular Ca²⁺ availability affects *S. aureus-induced* Ca²⁺ signalling in macrophages. However, in cells infected with the Δ*esxC* mutant, Ca^+2^ peaks occurred much earlier and were lower in magnitude [median 1.15 (IQR 1.07 - 1.6) hpi)], and the presence of EGTA did not impact the calcium signalling any further (Figure 6 B, E-H). These data indicate that EsxC interferes in early Ca²⁺ activation and amplification of the Ca²⁺ flux.

Mechanistically, these findings indicate that the T7SS effector EsxC, regulates the timing of Ca²⁺ signalling in macrophages. Together, the data suggest that T7SS-dependent effector secretion shapes host Ca²⁺ dynamics by suppressing early Ca²⁺ signalling events, which may underlie the delayed initiation of necroptosis and pyroptosis.

### Staphylococcal T7SS proteins affect skin infection outcomes

The T7SS is known to contribute to virulence in a range of murine infection models^32,36,42,44,67–69^. Given the effects that we observed for T7SS and its effectors EsxA and EsxC on macrophage cell death and our previous findings on epithelial cell death, we studied T7SS mutants during localised infection utilising a skin-abscess mouse model. At 24 hpi, Δ*essC*-infected mice had significantly decreased colony counts from the skin abscess tissue compared to WT-infected but not at 72 hpi (Figure 7A). Additionally, Δ*essC*-infected mice had significantly decreased abscess area compared to WT-infected (Figure 7B). However, Δ*esxC*-infected mice displayed no decrease but had a significantly increased bacterial burden in abscesses compared to WT-infected mice, with no difference in abscess area between groups (Figure 7C, D). Interestingly, Δ*esxA*-infected mice also showed a slight, statistically non-significant increase in abscess colony counts compared to WT-infected mice, but this was accompanied by an increased abscess area and abscess scores, with increased abscess sizes at 96 hpi (Figure 7E-H). The increased abscess size could indicate an altered immune response associated with the lack of EsxA, which also affects secretion of other effectors. The distinct responses elicited by T7 effector mutants may point towards effector-specific responses during infection.

**Figure 7.**
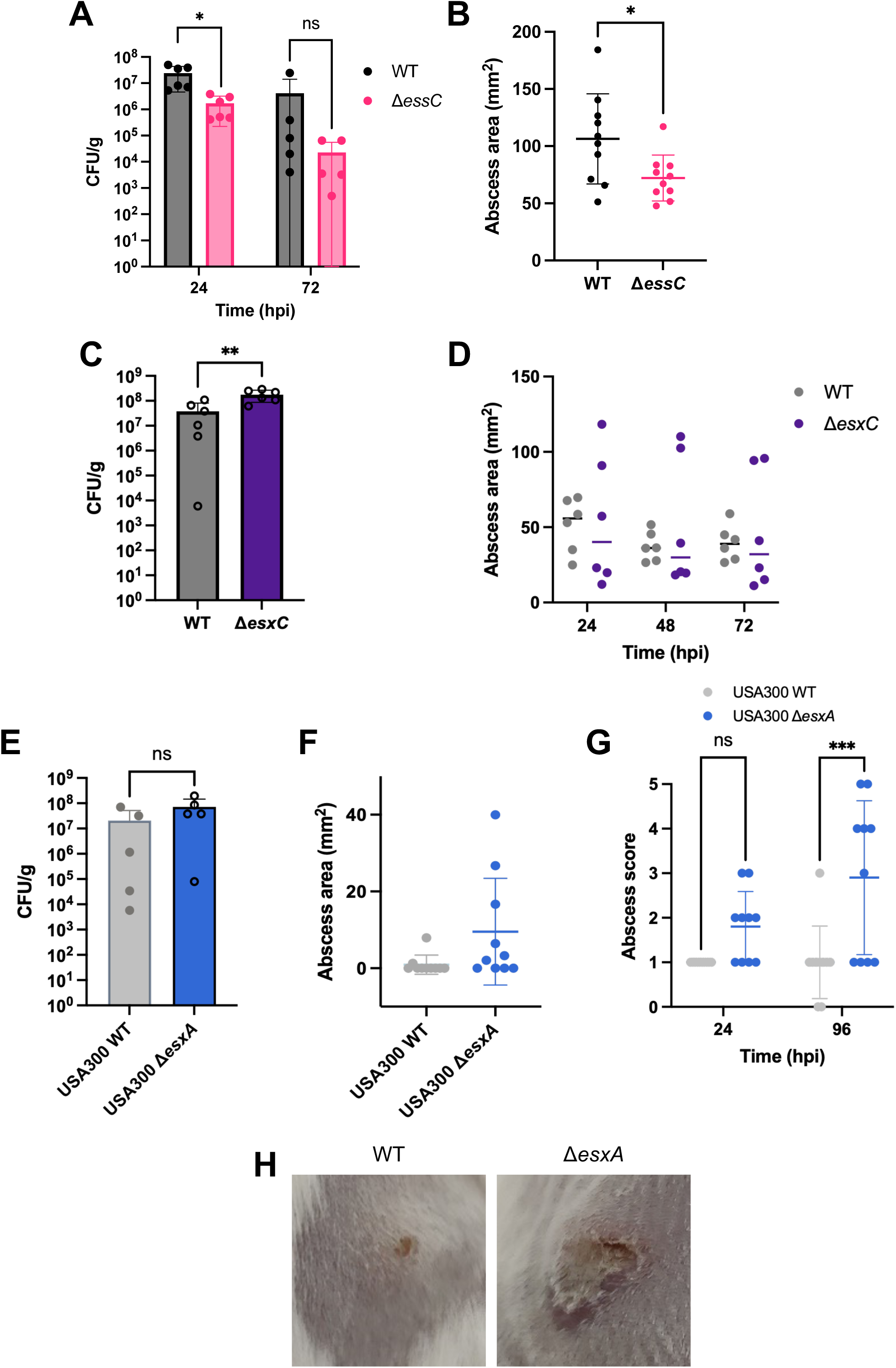
The staphylococcal T7SS impacts skin infection. Mice were infected on both flanks with JE2 or USA300 *S. aureus* WT and mutant strains as described in Methods. **A)** JE2 WT and Δ*essC* enumerated in the skin by colony counts at 24 and 72 hpi, n = 6/group, 2 lesions/mouse, mean ± SD, two-way ANOVA with Šídák’s multiple comparisons test, *P < 0.05. **B)** Abscess area measured in JE2 WT-and Δ*essC*-infected mice at 24 hpi, unpaired t-test, *P < 0.05. **C)** JE2 WT and Δ*esxC* enumerated in the skin by colony counts at 72 hpi, n = 3/group, 2 lesions/mouse, mean ± SD, shown, statistical significance calculated by unpaired t-test, **P < 0.01. **D)** Abscess area measured at 24, 48, and 72 hpi. Differences were not statistically significant by a 2-way ANOVA. **E)** USA300 WT and Δ*esxA* enumerated in the skin by colony counts at 96 hpi, n = 5/group, 2 lesions/mouse, mean ± SD, Differences were ns by a Mann-Whitney test. **F**) Abscess area of USA300 WT- and Δ*esxA*-infected mice at 96 hpi, unpaired t-test between strains **P < 0.01. **G)** Abscess score was calculated based on abscess morphology where 0 = clean, 1 = bump, 2 = white patch, 3 = white patch with pink/red external, 4 = white patch with pink/red external and brown centre, and 5 = open wound/open abscess. 2-way ANOVA with Šídák’s multiple comparisons test, ***P < 0.001. **H)** Representative images of infected skin abscesses at 96 hpi from WTaand Δ*esxA*-infected mice.

## Discussion

Macrophages are central to controlling bacterial infections by their ability to kill and present antigens to the adaptive immune system. However, they also provide a key intracellular niche for bacterial persistence for many pathogens including *S. aureus*. We report here for the first time how the staphylococcal type VII bacterial system (T7SS) proteins, which are strongly associated with bacterial virulence, play a key role in manipulating macrophage cell death. Our data show that the T7SS delays the formation of pore-induced trap (PIT) structures during macrophage infection, subsequently delaying macrophage lysis and bacterial escape. We demonstrate that at a molecular level, T7SS delayed the induction of the inflammatory cell death pathways, necroptosis and pyroptosis. Furthermore, the T7SS effector EsxC alters the timing and magnitude of calcium signalling in macrophages, which likely impacts the initiation of cell death pathways. T7SS-mediated modulation of cell death resulted in wider changes in chemokine expression and efferocytosis of infected macrophages, factors that may contribute to bacterial survival within tissues during infection (Figure 8).

**Figure 8.**
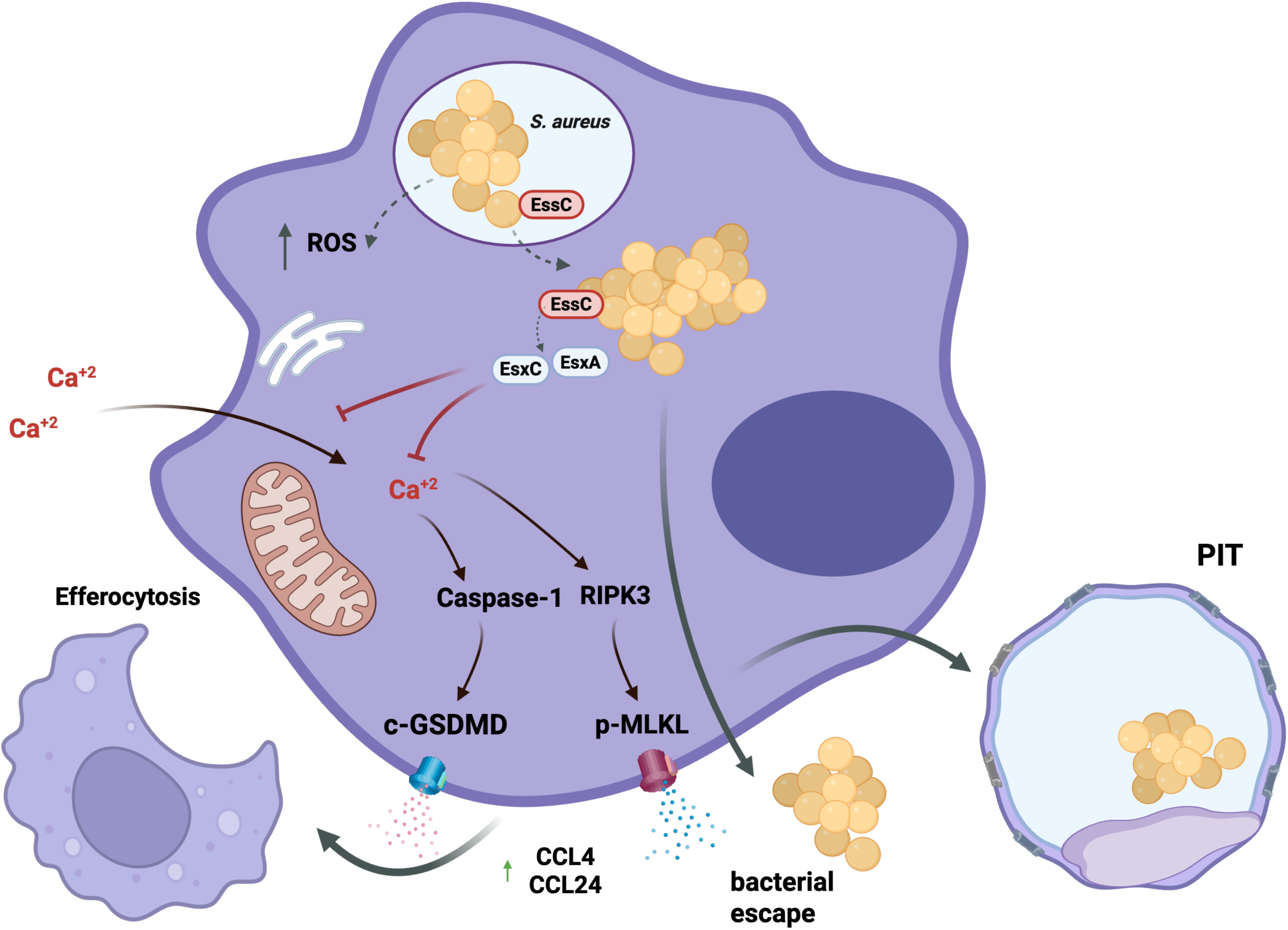
Model for the role of T7SS during macrophage infection. A proposed model for staphylococcal T7SS function during intracellular macrophage infection. Following phagocytic uptake, *S. aureus* survives and replicates intracellularly, encountering oxidative stress and triggering a rise in cytosolic Ca^2+^. T7SS effectors, including EsxA and EsxC, are exported by the transporter EssC and are presumed to be delivered into the cytosol. EsxC modulates the timing of intracellular Ca^2+^ signalling, delaying activation of Caspase-1 and RIPK3 signalling, resulting in delayed pyroptosis and necroptosis pathway activation, respectively. This eventually leads to delayed macrophage PIT formation and/or membrane disruption and subsequent bacterial release. The T7SS also alters chemokine expression and promotes efferocytosis of infected macrophages by neighbouring cells, which may aid in bacterial dissemination. Dashed arrows indicate proposed links

Previous research has hinted that PIT-like structures that balloon via membrane pores were observed upon induction of necroptosis and pyroptosis pathways^70,71^. Davis *et al*., 2019 describe “pyroptotic bodies” which lose their mechanical resilience; interestingly, all major cytoskeleton components were disrupted, similar to the complete loss of F-actin observed here during PIT formation^72^. LDH release was also noted to occur without rupture which may explain divergences between cytotoxicity assays and morphology kinetics observed here. We believe this is the first report of PIT structures during intracellular *S. aureus* infection. PITs have previously been described to entrap *Salmonella typhimurium* and *Listeria monocytogenes in vitro* as a result of inflammatory cell death^47,51^, and have been suggested to aid in the clearance of entrapped pathogens by secondary phagocytes^47^. However, during *S. typhimurium* macrophage infection, PITs exert a protective function, shielding the bacteria against neutrophil respiratory bursts and subsequent bacterial clearance^51^. While our data suggest a role for the T7SS in PIT formation, the relevance of PIT formation during *S. aureus* infection and staphylococcal factors that trigger PIT formation require further study. Similar to other pathogens, *S. aureus* containing PITs are likely efferocytosed during infection, as we demonstrate in infection assays *in vitro*. As these structures were efferocytosed by secondary macrophages in this and in previous studies^47,51^, we propose that efferocytosis could be expanded to include the clearance of such inflammatory corpses and termed “effero-PIT-osis”. Interestingly, despite Δ*essC*-infected macrophages forming more PITs, they were efferocytosed less often compared to WT-infected cells. As Δ*essC*-infected cells lysed earlier and released bacteria to seed microcolonies, PITs may rupture before they are engulfed, shortening the window for uptake in our assays. The T7SS-dependent differences in chemokine production in addition to signals released from dying cells could alter recruitment or recognition by neighbouring macrophages. Although the efferocytosis of PITs is not T7SS-dependent, T7SS promotes efferocytosis of infected cells, suggesting that it may aid *S. aureus* phagocytosis into new macrophages, potentially enabling bacterial dissemination from the site of infection.

*S. aureus* is known to induce both forms of programmed necrosis, necroptosis and pyroptosis in macrophages^19,21^. Multiple inflammatory pathways are likely triggered by staphylococcal factors like the alpha haemolysin, leading to cell death morphologies arising from multi-pathway executioner pores such as MLKL and GSDMD at the same time^73,74^. Here, we report T7SS-mediated delay of the induction of these cell death pathways during early macrophage infection. It is perhaps not surprising the T7SS affects multiple pathways as intricate crosstalk exists between the programmed cell death pathways. For example, activation of necroptosis could lead to pyroptosis induction via the NLRP3-Caspase-1 axis which is promoted by RIPK3 and MLKL activation and results in IL-1β processing^75,76^. Given that multiple pathways are activated by *S. aureus*, a PANoptosome, a supramolecular complex, containing various cell death pathway proteins, including RIPK1 and RIPK3, and Caspases-1, and -8, which work in synergy to execute cell death may be involved^77^. Additionally, in immune primed macrophages, there may be a redundancy between the inflammatory pathways. In GSDMD^-/-^ mice, where initial cell death is diminished during viral infection, inflammatory death was seen to increase later due to RIPK3 and Caspase-8 activation^78^.

Interestingly, the transcriptional up-regulation of oxidative stress genes from intracellular Δ*essC* suggests that T7SS may have an impact on oxidative stress during intracellular infection and aligns with previous observations in planktonically cultured T7SS mutants^39,40^. Moreover, the T7SS has previously been implicated in staphylococcal iron homeostasis^41^ which may in turn affect immune pathways and cell death. However, there was no significant impact of the T7SS on the induction of ferroptosis, another form of programmed cell death associated with iron-dependent lipid peroxidation and oxidative stress^79^.

Previous studies from our group and others have demonstrated that T7SS effector EsxA delays apoptotic death in epithelial and dendritic cells^30,31^, although the signalling mechanisms involved in this delay remain unclear. Recently, the T7SS effector EsxB was shown to reduce macrophage inflammation by directly inhibiting STING signalling^69^ which aligns with findings in this study. Unlike epithelial cells, in macrophages, we observe a T7SS-mediated delay with the inflammatory cell death pathways during intracellular infection. As T7SS effectors including EsxA and EsxC, are known to be co-dependent on other effectors for secretion^32,35,42^ we demonstrated direct effects on cell death by directly transfecting macrophages with selected effectors. However, it is important to note that studying effectors in isolation, in the absence of infection has its limitations, as the bacterium triggers cell death during infection, and moreover, different effectors could work together intracellularly to mediate the effects observed. While we have shown that EsxA and EsxC play likely roles in interfering with macrophage cell death, other effectors like EsaD, which are also induced intracellularly, are yet to be investigated and may also contribute to the timing of cell death. It is interesting that the staphylococcal T7SSb, in particular the effector EsxA, appears to function differently in macrophages compared to the well-studied counterparts in mycobacteria, where the T7SS (ESX-1) has been demonstrated to mediate phagolysosomal lysis, autophagy and cell death, and induction of type I interferons^80–83^. A recent study also showed that the Group B streptococcus EsxA can induce cytotoxicity in brain endothelial cells, and form pores on lipid membranes^84^. Our findings highlight that these specialised secretory systems have distinct functions that depend on the pathogen, cell type and the phase of infection.

Calcium has been strongly linked to cell death, with an overload of calcium triggering steps in the necroptosis pathway like RIP3 phosphorylation, or causing inflammasome and caspase activation, leading to pyroptosis^65,66,85^. Intracellular *S. aureus* can cause an increase in cytosolic calcium upon *S. aureus* infection, which involves an influx of Ca^2+^, from the extracellular environment and the endoplasmic reticulum, leading to the activation of calpains and caspases, triggering cell death in HeLa cells^8^. The cysteine protease staphopain A was implicated in the calcium induced cell death in this study^8^. In macrophages, however, blocking intra- or extracellular calcium affected autophagy and phago-lysosome fusion pathways, resulting in an increased survival of *S. aureus*^27,86^. Our data indicate that the T7SS may suppress calcium influx into the cell, at least from the extracellular space. Interestingly, only EsxC appears to modulate calcium homeostasis in our assays, which is aligned with recent findings that calcium-dependent effects of EsxC on the bacterial cell membrane and daptomycin sensitivity^41^. EsxC may modulate early Ca²⁺ release through influencing host Ca²⁺ channel engagement or interacting with calcium binding proteins, resulting in a delayed onset of calcium peaks during staphylococcal infection in macrophages. It is worth noting that while there were few transcriptional changes in calcium signalling-associated genes, ORAI-3, a Ca^2+^ channel protein which mediates store operated calcium channel activity, is significantly downregulated in the WT – but not in the Δ*essC*-mutant infected when compared to uninfected macrophages (Table S4). However, further investigations are necessary to understand how T7SS modulates calcium signalling pathways. Indeed, other non-calcium dependent pathways may also contribute to the suppression of cell death elicited by EsxA and potentially other T7SS effectors.

The precise functions of the staphylococcal T7SS during infection remains unclear – although studies suggest a role in immune modulation^30,43,44,67,69,87^. During bloodstream infections, the T7SS protein essE (esaE), which mediates EsaD secretion, was shown to induce IL-12 and IL1-β, and suppress production of RANTES (CCL5) from macrophages^43^. Our data show an induction of CCL3, a CC chemokine; both CCL3 and CCL5 activate the same receptor CCR5, expressed by myeloid cells, targeted by staphylococcal pore forming leukocidin LukED^88^. The T7SS protein EsxB was also reported to modulate inflammatory cytokines produced by dendritic cells, which in turn impacted the T-cell response, in particular IL-17 producing T-cells^30^, and its interaction with STING was further demonstrated to suppress the induction of proinflammatory cytokines during murine infection^69^. It is interesting that we see distinct effects on the size of the skin abscess and in bacterial survival for Δ*essC* (where all secreted effectors are absent) versus Δ*esxA* and Δ*esxC*, where presumably certain co-dependent effectors are absent. This indicates that while the entire system is necessary for bacterial survival in the host, individual effectors modulate the local immune responses which control the size of the abscess. A potential explanation for the differences between Δ*esxA* and Δ*esxC* phenotypes is that although both EsxA and EsxC protect against macrophage cell death and are reported to interact with each other *in vitro*, it is possible that secretion is modulated differently during infection or that they may be associated with distinct sets of bacterial effectors or host proteins inside macrophages. Further studies on effector expression, the abscess architecture and immune cell composition, along with inflammatory cytokine markers are necessary with these and other effector mutants to fully understand the contribution of individual T7SS effectors in staphylococcal skin abscesses.

In conclusion, we report here how *S. aureus* uses the T7SS to slow down the macrophage cell death programmes that are triggered in response to this pathogen, enabling the creation of an early niche to multiply and later escape from cells, or get efferocytosed by other macrophages, aiding dissemination. We show that the T7SS suppresses calcium signalling within cells, which delays activation of inflammatory pathways, subsequent changes to chemokine responses that may ultimately impact immune recruitment and clearance. This study highlights the importance of this intriguing group of proteins in immune modulation, further supporting the development of T7SS effectors as vaccine candidates.

## Acknowledgements

We would like to thank Medical Research Council for funding this study (grants MR/N010140/1 and MR/X00161X/1). Richard D. Allen and Kate E Watkins were funded by the University of Warwick MRC-doctoral training programme.

## Author contributions

R.D.A, RF, G.C, and P.A. performed the experiments for this study.

R.D.A, K.E.W, RF, M.U. were involved in designing the experiments in this study.

R.D.A, and M.U. wrote the manuscript. All authors were involved in reviewing the manuscript. We declare no conflicts of interest.

## Materials and Methods

### Bacterial strains and culture conditions

The bacterial strains used in this study are listed in Table S1. Primers used during cloning are listed in Table S2.

#### Macrophage-*S. aureus* intracellular infection

THP-1 macrophages (differentiated as described in Supplementary methods) were seeded at a density of 2.5 x 10^5^ cells/ml for infection and time-lapse assays, and at 5 x 10^5^ cells/ml for immunoblot and RNA-seq assays. BMDMs were seeded at a density of 1x10^5^ cells/ml for time-lapse assays, and at 2 x 10^5^ cells/ml for infection assays. *S. aureus* strains were cultured overnight and then subcultured to OD 1.5 (∼5 x 10^8^/ml), whereby strains were prepared for infection by dilution into RPMI+10% FBS to achieve an MOI of 10:1 (bacteria: macrophage). THP-1 macrophages and BMDMs were pre-treated with 10 μM necrosulfonamide (NSA) (Merck, 480073) or 20 μM VX-765 (Invivogen) for at least 1 hr prior to infection where appropriate. Macrophages were infected at an MOI of 10:1 (bacteria: macrophage) for 1 hr before extracellular killing of bacteria with 20 μg/ml lysostaphin (Sigma), and 50 μg/ml gentamicin. Macrophages were then resuspended in RPMI + 10% FBS without antibiotics until appropriate timepoint for lysis to measure internalisation of bacteria, microscopy, or immuno-assays.

#### Time-lapse microscopy

Infections were carried out as previously described in 4-well Permanox plastic chamber slides (Nunc Lab-Tek, ThermoFisher Scientific) with YFP-expressing *S. aureus* strains grown in chloramphenicol 10 μg/ml. After initial killing of extracellular bacteria with gentamicin and lysostaphin, cells were washed with PBS and refreshed in RPMI +10% FBS containing no antibiotics. Fluorescent and phase contrast images were taken every 10 min for 3 fields of view in each well from 2 hrs to 24 hrs post-infection. Images were acquired using a Leica DMi8 widefield fluorescent microscope with CO_2_ stage top incubator (Okolab). Quantification of time-lapse microscopy was carried out blinded by manually tracking the morphological changes of 50 infected cells in each field of view at various time points. Morphological features include spindle-shaped (elongated cell body with apical cytoplasmic protrusions), fried egg-shaped (enlarged ameboid cell body), rounded (smaller round cells), and PIT-like structures (immobilised, ballooned circular structure, condensed organelles), as previously described^47^. For BMDM infections, macrophage-macrophage interactions were also noted, including efferocytosis.

#### Lipid staining of PITs

Following manufacturer’s instructions, HCS LipidTOX^TM^ Red Neutral Lipid Stain (Invitrogen) was used to stain lipid granules within fixed samples of THP-1 macrophages uninfected and infected with WT JE2 *S. aureus*. Imaging of stained samples carried out on Cytation5 (Agilent BioTek), Leica BMi8 inverted widefield fluorescent microscope with ORCA-Flash4.0 V2 digital CMOS camera (Hamamatsu Photonics), and UltraVIEW VoX (PerkinElmer).

#### Bacterial escape and microcolony formation

THP-1 macrophages, BMDMs, and iBMDMs were challenged by JE2 *S. aureus* as described above without antibiotic incubation after extracellular killing. For THP-1 macrophages, supernatants were plated for CFU counts at 6 and 10 hpi to assess bacterial escape, subsequently macrophages were lysed to obtain cell-associated CFU/ml. Macrophage microcolonies were stained with LIVE/DEAD BacLight Bacterial Viability Kit (ThermoFisher Scientific) following manufacturer’s instructions.

#### Efferocytosis assay

iBMDMs were seeded at 1 x 10^5^ cells/well in μ-Slide 4 Well Ph+ chamber slides (Ibidi) and separate iBMDM populations were stained with CellTracker^TM^ (Invitrogen) CMPTX at 1 μM or CMAC at 5 μM for 20 min following manufacturer’s instructions. CMPTX-stained iBMDMs were then infected as described above and refreshed after antibiotic clearance in RPMI+10% FBS containing CMAC-stained iBMDMs that were removed by Accutase^TM^ treatment, at a 1:1 ratio. Interactions between populations were imaged live on a Zeiss LSM900 with LSM Plus processing, every 10 min for 24 hrs.

#### Cell death assays

CytoTox 96® Non-Radioactive Cytotoxicity Assay (Promega) was used following manufacturer’s instructions to determine % cytotoxicity during *S. aureus* challenge of THP-1 macrophages as previously described, at 2, 4, 6, 8, and 10 hpi.

Caspase-1 activation in THP-1 macrophages during *S. aureus* challenge as previously described was assessed using Caspase-1 Glo Assay (Promega) following manufacturer’s instructions at 2 hpi.

#### Protein lysates and immunoblotting

At the appropriate time-points post infection, cells were treated with radioimmunoprecipitation (RIPA) buffer (Fisher Scientific) containing 1x Halt protease and phosphatase single-use inhibitor cocktail (Fisher Scientific, 10127963) on ice. Lysates were centrifuged to remove cellular debris. 20-25 μg of protein were run on 15% Mini-PROTEAN TGX Protein gels (BioRad) and electrotransfered using Trans-Blot Turbo Transfer System (BioRad) with Mini PVDF transfer packs (BioRad). Blots were blocked with 5% BSA (Millipore) in 1X TBST (Tris-buffered saline plus Tween20). Immunodetection was carried out with rabbit anti-phosphorylated MLKL ser358 (p-MLKL) [Cell Signalling Technology (CST), 91689], rabbit anti-MLKL(D216N) (CST 14993S), rabbit anti-phosphorylated RIPK3 (p-RIPK3) (CST 54105), rabbit anti-RIPK3 (CST 54105), rabbit anti-cleaved GSDMD (CST 36425S), rabbit anti-GSDMD (Abcam, EPR19828), rabbit anti-NLRP3 (CST, 15101S), rabbit anti-Caspase 4 (CST4450), rabbit anti-GPX4 (CST 52455), mouse anti-β-actin (Sigma Aldrich A5316), mouse anti-α-tubulin (Sigma Aldrich T5168). Blots were developed with Pierce horseradish peroxidase (HRP) Western blot substrate (ThermoFisher Scientific). Band intensities were quantified using ImageJ and relative densities were calculated by normalisation to either β-actin or α-tubulin bands using ImageJ (version 2.14.0/1.5f).

#### Dual RNA-seq analysis

*S. aureus* challenge of THP-1 macrophages was carried out as described above and incubated without antibiotic after extracellular killing for 2 and 6 hpi. At the appropriate timepoint, cells were washed with PBS, before the addition of 700 μl LETs buffer (0.1 M LiCl, 0.01 M Na2EDTA, 0.01 M Tris-Cl pH 7.4, 0.2% SDS) and subsequently frozen at -80°C. Total RNA was extracted from the samples by bead-beating with lysing matrix B beads (Invitrogen), addition of TRIzol Reagent (ThermoFisher Scientific) and chloroform. RNA was precipitated with isopropanol, centrifuged, and washed with 75% ethanol and resuspended in RNAse-& DNAse-free water. DNA was removed via TURBO DNA-free Kit (Invitrogen) following manufacturer’s instructions, and samples were cleaned up with LiCl Precipitation Solution 7.5 M (Invitrogen). RNA integrity was assessed via Agilent 6000 RNA Pico (Agilent Technologies) and RNA purity and contamination checked via NanoDrop One (ThermoFisher Scientific). Subsequently, rRNA-depletion, library preparation, and Illumina sequencing was carried out by GeneWiz (Azenta). Bacterial samples were sequenced to a depth of at least 10 million reads, human samples to a depth of at least 40 million reads, and mixed samples to a depth of at least 60 million reads. FASTQ sequencing files were initially assessed by FastQC^89^ and followed by trimming of adapter contamination using Trimmomatic tool (version 0.39)^90^. Trimmed reads were aligned by Hierarchical Indexing for Spliced Alignment of Transcripts v2.1.0 (HISAT2)^91^, to individual or concatenated genomes: human genome GRCh38.p14 (release 46) with comprehensive gene annotation (GENCODE), and *S. aureus* JE2 genome (NCBI Reference Sequence GCF_002085525.1).

Mapping analysis and file conversion was carried out using Bowtie 2 and SAMtools. Gene expression counts were generated by LiBiNorm^92^. DEGs were generated by DeSEq2 (version 1.42.1), in RStudio (version 2023.06.1+524), where DEGs were considered significantly differentially expressed with adjusted p-values below 0.05 and log_2_ fold change ≥1 and ≤-1. Poorly annotated genes were further investigated by BLASTX and BLASTP for homology. Heatmaps were generated using the R “pheatmap” package and normalised counts, and non-biased hierarchical clustered heatmaps were generated through elbow plots where the complete linkage method and Euclidean distance, followed by K-means clustering were used to determine the optimal number of clusters.

All sequencing reads were submitted to the EMBL-European Bioinformatics Institute, Array Express (Accession number E-MTAB-16250).

#### Macrophage transfection and flow cytometry

*esxA* and *esxC* were individually cloned by amplification from JE2 WT gDNA into vector pEGFP-N1, containing constitutively expressed eGFP. iBMDMs were transfected by nucleofection using Lonza Mouse Macrophage Nucleofector^TM^ Kit (Lonza) following manufacturer’s instructions. At 12 hr post-transfection, adherent macrophages were dislodged with Accutase (Corning) and stained with Annexin V Blue Pacific (Invitrogen) and 0.2 mg/ml propidium iodide (PI), then fixed with 4% PFA and kept on ice until analysis via flow cytometry on LSRFortessa^TM^ (BD Biosciences). For a positive control, eGFP-transfected cells were treated with staurosporine 5 mM for 30 min prior to dislodgment and staining. Data was collected from a minimum of 20,000 macrophages and 3,000 GFP+ macrophages using BD FACSDiva software (BD Biosciences) and analysed using floredia.io online tool (https://floreada.io) and plots generated on FCS Express 7 version 7.28.0019 (De Novo Software). For gating, macrophages were selected from forward and side scatter of unstained samples and subsequently for stained samples, eGFP+ cells were gated from the macrophage cells. eGFP+ cells were further gated for PI+ and AV+, for which the percentage of PI-AV+ cells were calculated for each sample condition.

#### Calcium flux assay

Intracellular Ca²⁺ signalling was assessed using Fluo-4 and Fluo-8 to measure both single-cell and population-level Ca²⁺amplitude, respectively. First, iBMDMs were seeded at 1 x 10^5^ cells/well in 8 Well Permanox plastic chamber slides (Nunc Lab-Tek, Thermo Fisher Scientific) overnight. iBMDMs were loaded with 5 µM Fluo-4-AM (Thermo Fisher Scientific) or Fluo-8-AM (Abcam) as per manufacturer instructions. For extracellular Ca²⁺ chelation, HBSS was supplemented into cell culture media with 3 mM EGTA before infection.

Macrophages were infected at MOI 10:1 with *S. aureus* strains expressing pMV158-mCherry. Time-lapse fluorescence imaging was performed at 5 min intervals for Fluo-4 (for a total of 200 min), and 1 min intervals for Fluo-8 (for a total of 120 min). Single cell image analysis (Fluo-4) was performed using Fiji; ROIs that correspond to individual macrophages were background-subtracted and normalised to the 0–20 min baseline to obtain relative fluorescence traces. The maximum amplitude was determined as the highest relative fluorescence value measured, and peak latency was defined as the time point of maximum amplitude. For Fluo-8 experiments, fluorescence [RFU (Max-Min)] values were background-subtracted and normalised to the baseline fluorescence (F₀). F₀ was defined as the mean fluorescence intensity at the initial time point prior to Ca²⁺ mobilisation. Changes in fluorescence at each time point were expressed as ΔF/F₀, where ΔF = F(t) − F₀, providing a normalised measure of cumulative Ca²⁺ mobilisation across the field of view. A total field percentage fluorescence was quantified at each time point and expressed as ΔF/F₀ × 100.

#### Murine skin abscess model

Female BALB/c mice, aged 7–8 weeks, were obtained from Charles River Laboratories, UK (license PP9171369). Animals were housed in individually ventilated cages under standardized conditions, including a 12-hour light/dark cycle. All mice were acclimatized at the University of Warwick Rodent Facility prior to the initiation of experiments. On the day of the experiment, mice were divided into three groups, and both flanks of all mice were shaved. Mice were injected intradermally with *S. aureus* (3-6 mice per group) or mutant strain (3-6 mice/ group) at 1 x 10^6^ CFU/mouse in 50 μl PBS or injected with PBS only on right and left flanks. Mice were monitored regularly for signs of distress or ill health and weighed daily. Abscesses on both flanks were measured and scored daily. At 24, 48, 72, and 96 hpi mice were sacrificed via cervical dislocation. Infected skin samples were weighed and homogenised in PBS using a FastPrep (MP Biomedicals), at 4 m/s for 60 seconds for a total of 6 cycles. Homogenates were diluted in PBS and CFU/ml was determined. CFU counts were normalized to lesion weight.

### Statistical analysis

All statistical analysis was carried out using GraphPad Prism version 9.5.1 or later for Mac (GraphPad Software) and subjected to a Shapiro-Wilk normality test to assess data distribution. For pairwise analysis on two groups of parametric data, Student’s paired or unpaired t-test was used, and for non-parametric data, a Mann Whitney test was used with Holm-Šídák’s multiple comparison test where appropriate. For pairwise analysis on multiple groups that were parametric, a two-way ANOVA was used followed by Dunnett’s multiple comparisons test, and for non-parametric data multiple paired t-tests were conducted with Holm-Šídák’s multiple comparisons or Kruskal-Wallis with Dunn’s corrections. For column analysis of multiple groups, a One-way ANOVA with Tukey’s multiple comparison test was used. Due to intra-replicate variation, fold change was calculated relative to the WT condition and assessed by one sample t-test against a hypothesised value of 1. For time-lapse analysis, a minimum of 150 cells per condition were quantified blinded across at least 3 biological replicates. Specific descriptions of analyses used are described in figure legends.

## Supplementary figure legends

**Figure S1. A)** *S. aureus* replicates and lyses the PIT structure to form a microcolony and **B**) intracellular *S. aureus* that do not replicate within PIT structures, scale bars represent 20 μm. **C)** LipidTOX staining of lipid granules and DAPI in uninfected and JE2 WT-infected THP-1 macrophages, with PIT-like structure enlarged, scale bars represent 20 μm. Quantification of uninfected macrophage morphologies during time-lapse microscopy for THP-1 macrophages (**D**) and BMDMs (**E**). Morphological fate of individual uninfected macrophages over time for THP-1 macrophages (**F**) and BMDMs (**G**). Data represents N = 3 for A, B, D-G and N = 2 and graphs display mean ± SD.

**Figure S2.** THP-1 macrophages were infected with JE2 WT and Δ*essC* for 1 hr at MOI 10:1 before extracellular killing. Cytotoxicity was measured by lactate dehydrogenase release across timepoints and presented as percentage. A 2-way ANOVA did not show any statistical differences between WT and Δ*essC* infected cells.

**Figure S3.** THP-1 macrophages were infected with YFP-expressing JE2 WT or Δ*essC* for 1 hr at MOI 10:1 before extracellular killing of bacteria and time-lapse microscopy conducted. Representative images of microcolony formation over time during infection of macrophages with WT or Δ*essC*. Scale bar represents 20 μm, and representative images from three biological independent experiments are shown.

**Figure S4.** Quantification of intracellular JE2 WT and Δ*essC* at 6 hpi during BMDM infection with and without necroptosis inhibitor NSA, presented as CFU/ml (**A**) and by relative fold change to WT (**B**), N = 3, mean ± SD, statistical significance was calculated by multiple paired t-tests with Holm-P*<0.05, P**<0.01. BMDMs were infected with JE2 WT with and without NSA inhibitor and assessed by timelapse microscopy. **C**) Quantification carried out blinded for PIT-like structures relative to WT-infected BMDMs, N = 3, mean ± SD, statistical significance calculated by one-sample t-test to WT, *P<0.05, **P<0.01.

**Figure S5.** THP-1 macrophages were infected with JE2 WT and Δ*essC* strains for 1 hr at MOI 10:1 before extracellular killing, with and without necroptosis inhibitor NSA. Quantification of cellular morphology blinded from time-lapse data: fried egg & spindle (black), PIT-like (pink), rounded (blue), and apoptotic-like (purple). Morphology presented for WT (**A**), WT + NSA (**B**), Δ*essC* (**C**), Δ*essC* + NSA (**D**), uninfected (**E**), and uninfected + NSA (**F**). N = 3 and graphs present mean ± SD. Quantification of cellular morphology blinded from time-lapse data. Quantification of the percentage of PIT (**G**) and apoptotic-like (**H**) morphologies during infection, N = 3, mean ± SD, statistical analysis by multiple paired t-tests with Holm-Šídák’s multiple comparison test for **G**) and a two-way ANOVA with Dunnett’s multiple comparisons test for (**H**) *P <0.05, **P < 0.01, and ***P <0.001.

**Figure S6.** A) Representative images of microcolonies stained with LIVE/DEAD in JE2 WT-infected THP-1 macrophages with and without NSA inhibitor at 10 hpi, scale bar represents 30 μm. **B**) Intracellular JE2 WT collected from 6 hpi and stained with LIVE/DEAD (SYTO9 in green and propidium iodide in red) and imaged at x100 magnification, scale bar represents 5 μm. **C**) THP-1 macrophages infected with JE2 WT and Δ*essC* for 6 hrs before being lysed to collect intracellular bacteria; lysed bacteria were used to infect a secondary set of THP-1 macrophages for 4 hrs, cells were lysed and CFU determined. Re-infection control represents number of bacteria from lysates of secondary set of THP-1 macrophages infected with an equivalent number of bacteria (to that collected after 6 hpi) from standard subculture. Mann-Whitney tests and Holm-Šídák’s multiple comparison test revealed statistical significance between 1^st^ and 2^nd^ infection for WT and Δ*essC*, presented as (*P<0.05, **P<0.01). Data represents N=3, mean ± SD, *P<0.05, **P<0.01.

**Figure S7** BMDMs were infected with JE2 WT, Δ*essC*, and Δ*essC*-C for 1 hr at MOI 10:1 and lysates were collected at 4 and 6 hpi. A) representative immunoblots probed with anti-NLRP3 and anti-α-tubulin loading control. B) quantification of NLRP3 bands normalised to loading control, presented as fold change relative to WT at each timepoint. N=3, mean ±SD and no statistical differences between WT and mutant groups.

**Figure S8.** Differentially expressed genes (DEGs) were generated from DESeq2 analysis of dual RNAseq counts. **A**) The number of DEGs from comparison of JE2 *S. aureus* from infected cells to planktonic control at each timepoint for WT and Δ*essC* at 2 and 6 hpi. **B**) The number of DEGs comparing WT-or Δ*essC*-infected macrophages to uninfected macrophages at 2 and 6 hpi. DEGs were selected under the threshold of p.adj < 0.05 and log2FC ≥1 and ≤-1. DEGs are represented at 2 hpi in blue, and at 6 hpi in orange. **C**) Leaf plots displaying the overlap of DEGs between WT and Δ*essC* condition, at 2 and 6 hpi for up- and down-regulated JE2 *S. aureus* and macrophage DEGs.

**Figure S9.** KEGG pathways enriched from Δ*essC*-vs WT-infected macrophage up- and down-regulated DEGs at 2 hpi (**A, C**) and 6 hpi (**B**). Number of DEGs identified in enriched pathway is represented by circle size (Count) and P-adjusted value (P. adjust) is represented by colour. KEGG pathway enrichment from up- and down-regulated WT-infected macrophage vs. uninfected macrophage DEGs at 2 hpi (**D, F**) and 6 hpi (**E**). Number of genes associated with the term is represented by circle size (count) and P-adjusted value is represented by colour.

**Figure S10. A**) Heatmap of cytokine expression JE2 WT or Δ*essC* infected THP-1 macrophages and uninfected macrophages, at 2 and 6 hpi. Red asterisks indicate significantly differential gene expression between Δ*essC* and WT when compared to uninfected macrophages. **B)** Normalised macrophage counts from dual RNA-seq showing key genes in the ferroptosis pathway. **C)** THP-1 lysates obtained at 6 hpi were probed with anti-GPX4 and with anti-α-tubulin (loading control). A representative immunoblot is shown. Quantification of GPX4 bands normalised to loading control is shown below the image and presented as fold change relative to WT. N = 3, mean ± SD.

**Figure S11 A**) Heatmap of normalised expression depicting transcript abundance of T7SS locus across bacterial samples. Expression is represented as a computed Z-score denoting high (red) and low (blue) expression across samples. **B**) Representative flow cytometry plots of eGFP-positive macrophages gated for Annexin V and PI (for data shown in Figure 5D) **C**) iBMDMs were transfected with either pEGFP-N1, pEGFP-N1+*esxA*, or pEGFP-N1+*esxC* via nucleofection. At 12 hrs post transfection, macrophages were stained with Annexin V (AV) Blue Pacific and propidium iodide (PI) and analysed by flow cytometry. Transfected cells were gated for eGFP expression. The number of PI+AV- and PI+AV+ macrophages relative to the empty control vector (eGFP) are shown. N = 3, mean ± SD and differences were not significant (ns) by a one-sample t-test.

**Figure S12.** Cumulative Fluo-8 fluorescence in macrophages following infection at MOI 10. Macrophages were loaded with Fluo-8 AM and infected with WT (USA300) or Δ*esxA.* Time-lapse fluorescence imaging was performed at 1 min intervals for 120 min. For each time point, fluorescence intensity was quantified across the entire field of view and expressed as cumulative % = (ΔF/F₀) × 100 relative to baseline. Data represents mean ± SEM, N =3.

**Table S1** Bacterial strains used in this study

**Table S2** Primers used in this study

**Table S3** Δ*essC*-infected vs. WT-infected macrophage DEGs

**Table S4** Unique DEGs for WT-or Δ*essC*-infected macrophages when compared to uninfected macrophages, and WT or Δ*essC* compared to planktonic

**Table S5** Δ*essC* vs. WT DEGs during intracellular infection

**Table S6** Bacterial iron and oxidative stress related DEGs during macrophage infection

**Table S7** Infected macrophage ferroptosis DEGs

