## Supplementary material for "The staphylococcal type VII secretion system delays inflammatory cell death during macrophage infection": Suppl figures S1-S12

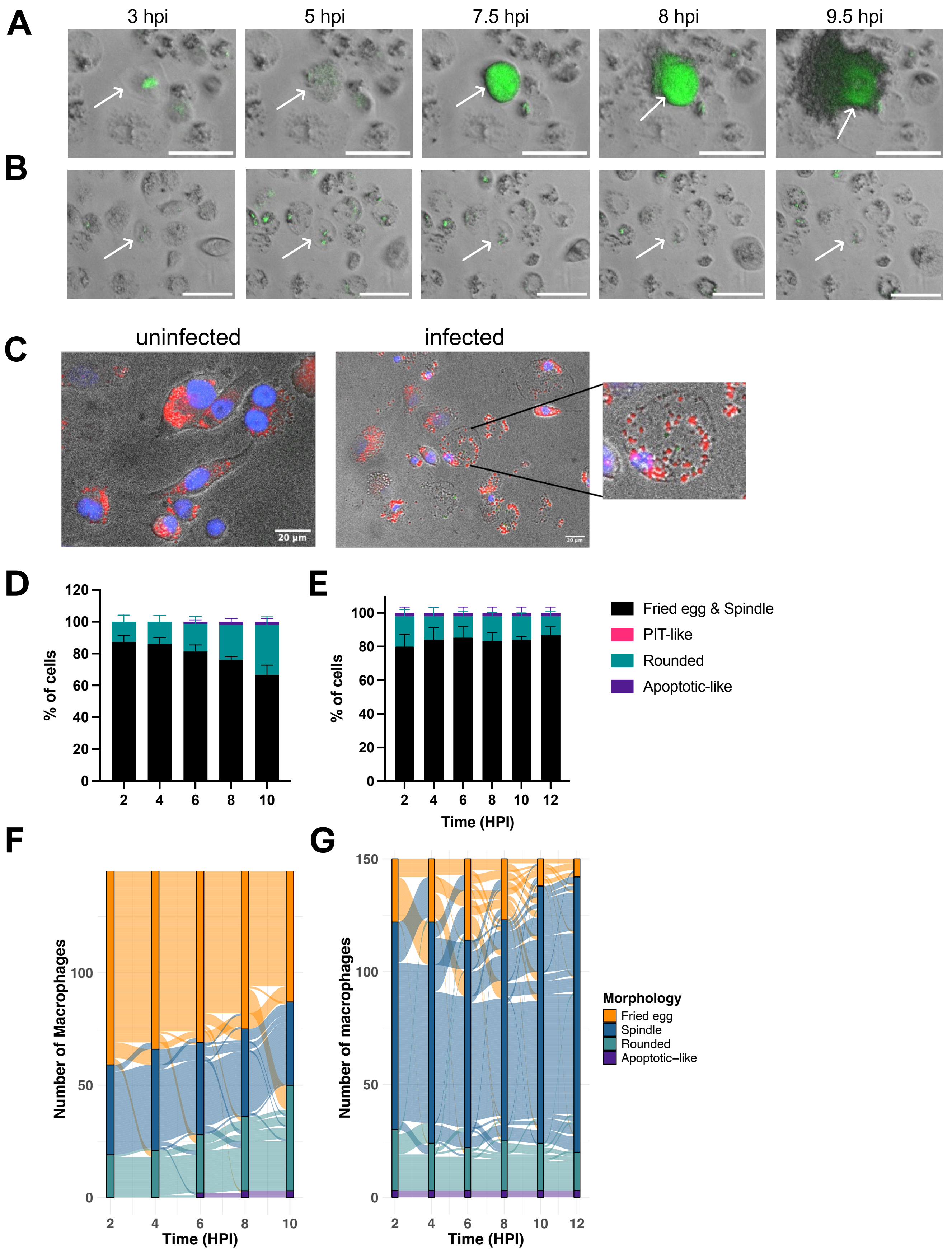


**Figure S1. A)** *S. aureus* replicates and lyses the PIT structure to form a microcolony and **B**) intracellular *S. aureus* that do not replicate within PIT structures, scale bars represent 20 µm. **C**) LipidTOX staining of lipid granules and DAPI in uninfected and JE2 WT-infected THP-1 macrophages, with PIT-like structure enlarged, scale bars represent 20 µm. Quantification of uninfected macrophage morphologies during time-lapse microscopy for THP-1 macrophages (**D**) and BMDMs (**E**). Morphological fate of individual uninfected macrophages overtime for THP-1 macrophages (**F**) and BMDMs (**G**). Data represents N = 3 for A, B, D-G and N = 2 and graphs display mean ± SD.

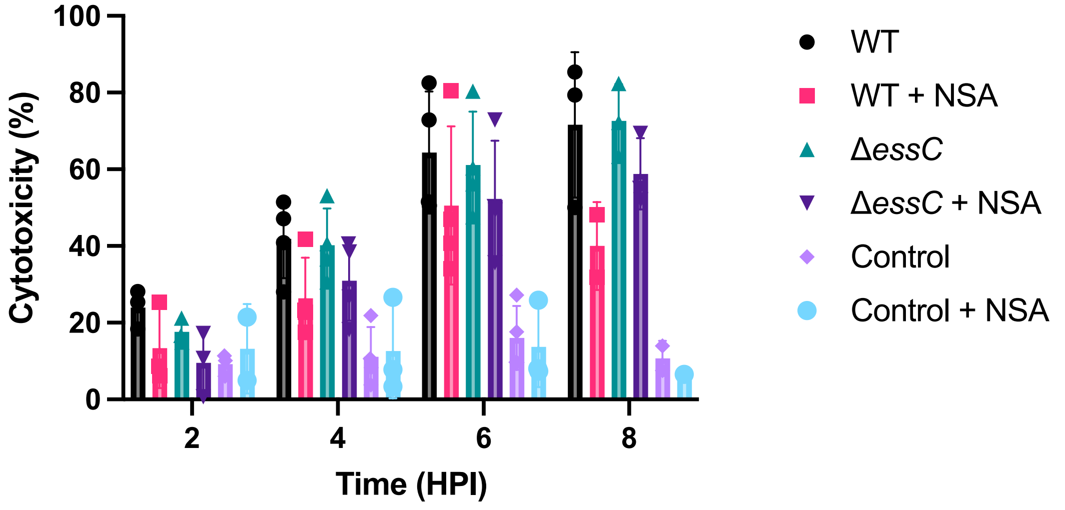


**Figure S2.** THP-1 macrophages were infected with JE2 WT and Δ*essC* for 1 hr at MOI 10:1 before extracellular killing. Cytotoxicity was measured by lactose dehydrogenase release across timepoints and presented as percentage. A 2-way ANOVA did not show any statistical differences between WT and Δ*essC* infected cells.

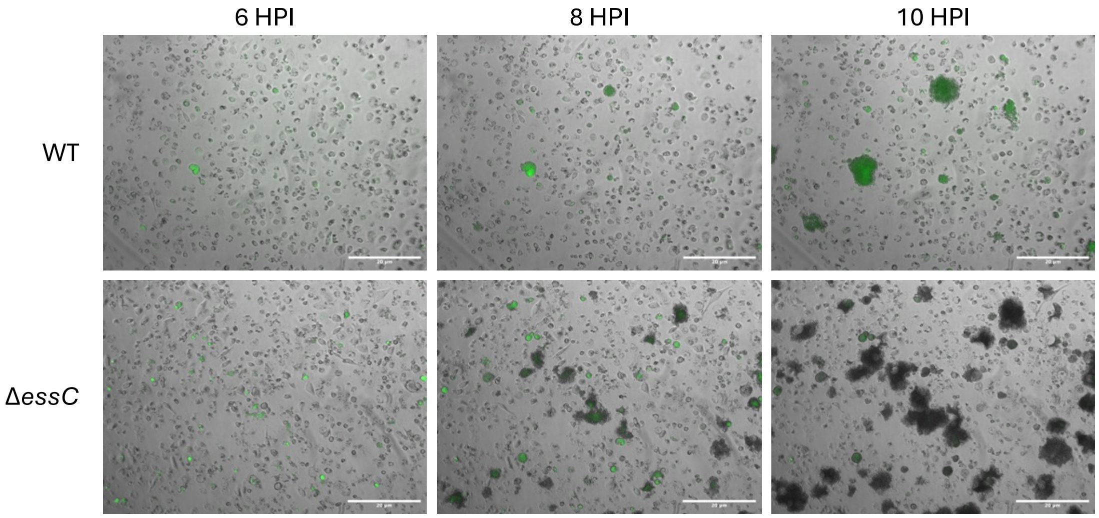


**Figure S3.** THP-1 macrophages were infected with YFP-expressing JE2 WT or Δ*essC* for 1 hr at MOI 10:1 before extracellular killing of bacteria and time-lapse microscopy conducted. Representative images of microcolony formation over time during infection of macrophages with WT or Δ*essC*. Scale bar represents 20 μm, and representative images from three biological independent experiments are shown.

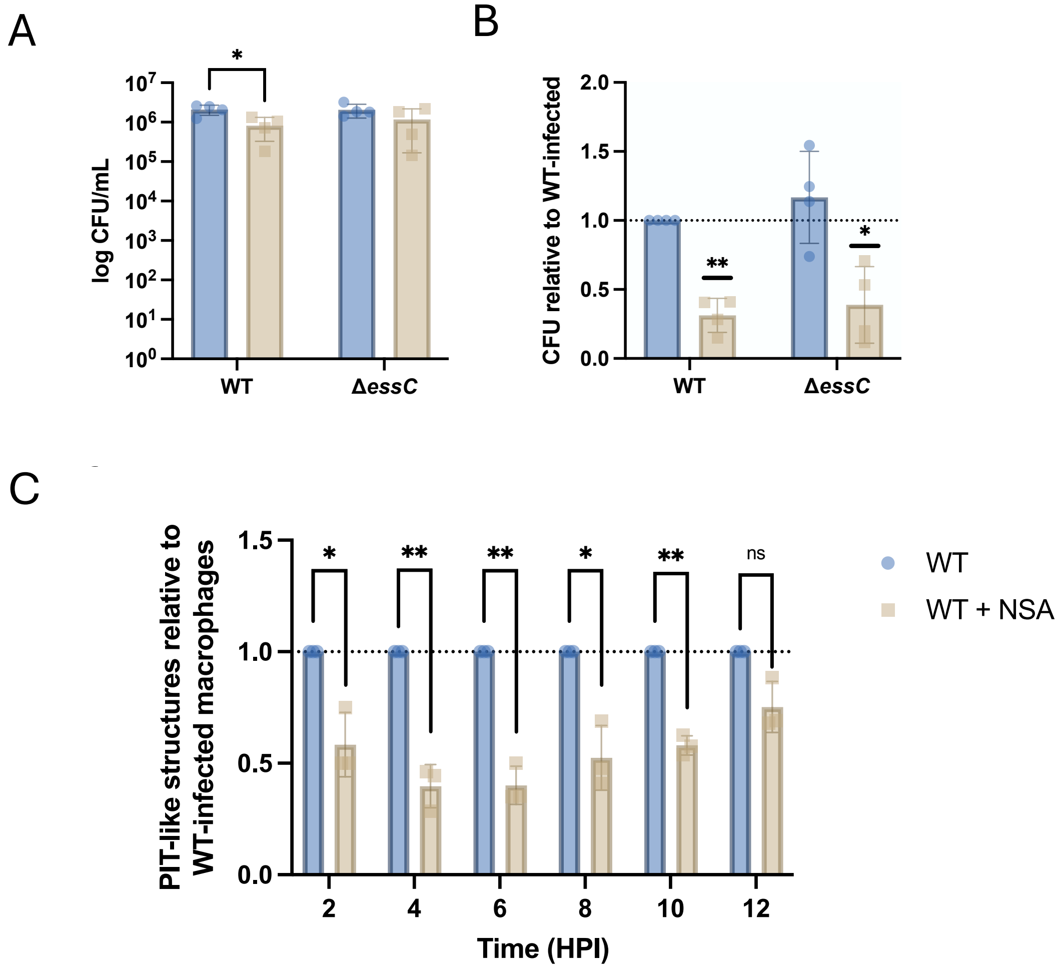


**Figure S4.** Quantification of intracellular JE2 WT and Δ*essC* at 6 hpi during BMDM infection with and without necroptosis inhibitor NSA, presented as CFU/ml (A) and by relative fold change to WT (B), statistical significance was calculated by multiple paired t-tests with Holm-Šídák's multiple comparisons tests for (A) and a one-sample t-test relative to WT for (B). BMDMs were infected with JE2 WT with and without NSA inhibitor and assessed by timelapse microscopy. C) Quantification carried out blinded for PIT-like structures relative to WT-infected BMDMs, statistical significance calculated by one-sample t-test to WT.

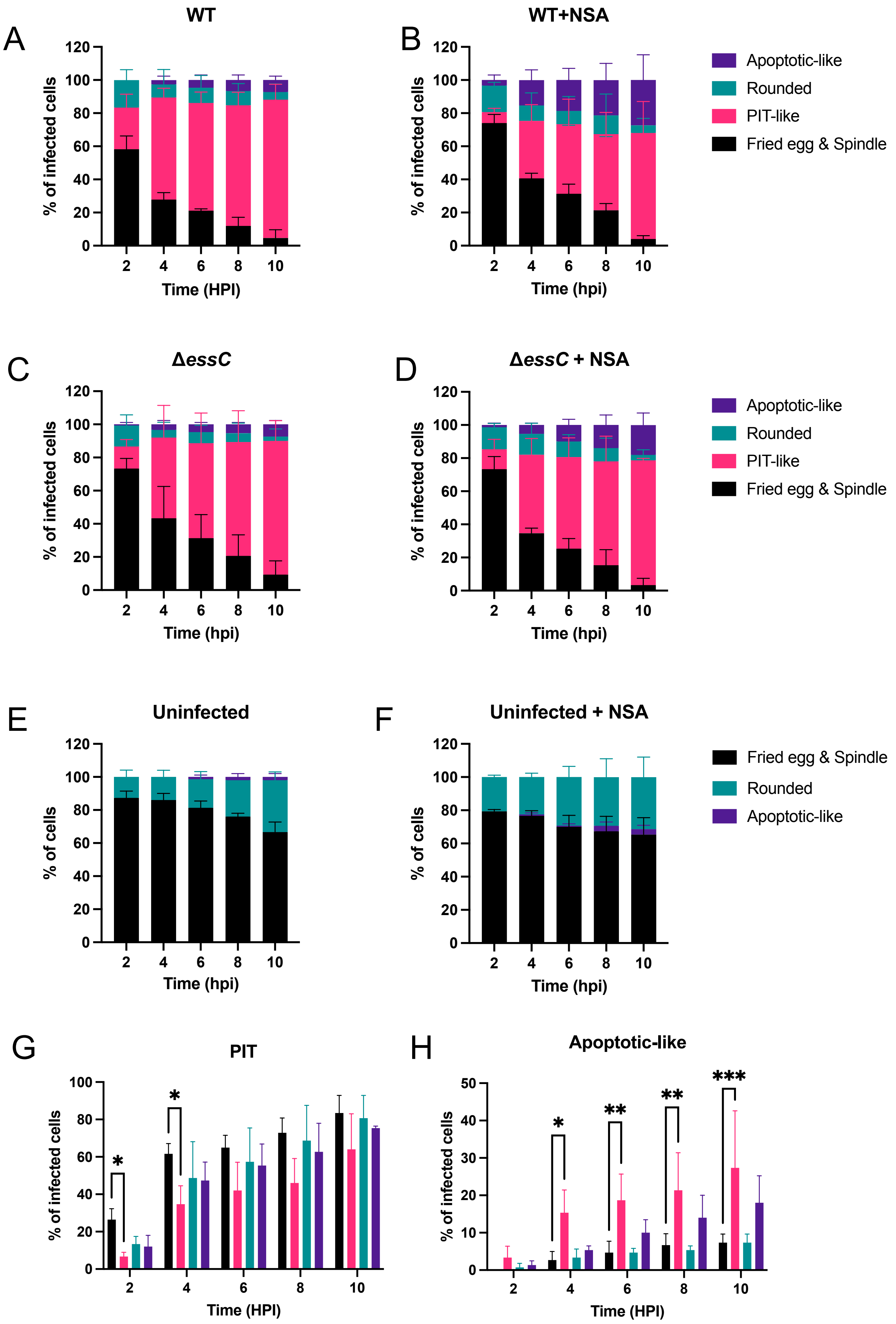
A
WT
% of infected cells
Time (HPI)
B
WT+NSA
% of infected cells
Time (hpi)
Apoptotic-like
Rounded
PIT-like
Fried egg & Spindle
C
ΔessC
% of infected cells
Time (hpi)
D
ΔessC + NSA
% of infected cells
Time (hpi)
Apoptotic-like
Rounded
PIT-like
Fried egg & Spindle
E
Uninfected
% of cells
Time (hpi)
F
Uninfected + NSA
% of cells
Time (hpi)
Fried egg & Spindle
Rounded
Apoptotic-like
G
PIT
% of infected cells
Time (HPI)
H
Apoptotic-like
% of infected cells
Time (HPI)

**Figure S5.** THP-1 macrophages were infected with JE2 WT and Δ*essC* strains for 1 hr at MOI 10:1 before extracellular killing, with and without necroptosis inhibitor NSA. Quantification of cellular morphology blinded from time-lapse data: fried egg & spindle (black), PIT-like (pink), rounded (blue), and apoptotic-like (purple). Morphology presented for WT (**A**), WT + NSA (**B**), Δ*essC* (**C**), Δ*essC* + NSA (**D**), uninfected (**E**), and uninfected + NSA (**F**). N = 3 and graphs present mean ± SD. Quantification of cellular morphology blinded from time-lapse data. Quantification of the percentage of PIT (**G**) and apoptotic-like (**H**) morphologies during infection, N = 3, mean ± SD, statistical analysis by multiple paired t-tests with Holm-Šídák's multiple comparison test for **G**) and a two-way ANOVA with Dunnett's multiple comparisons test for (**H**) *P <0.05, **P < 0.01, and ***P <0.001.

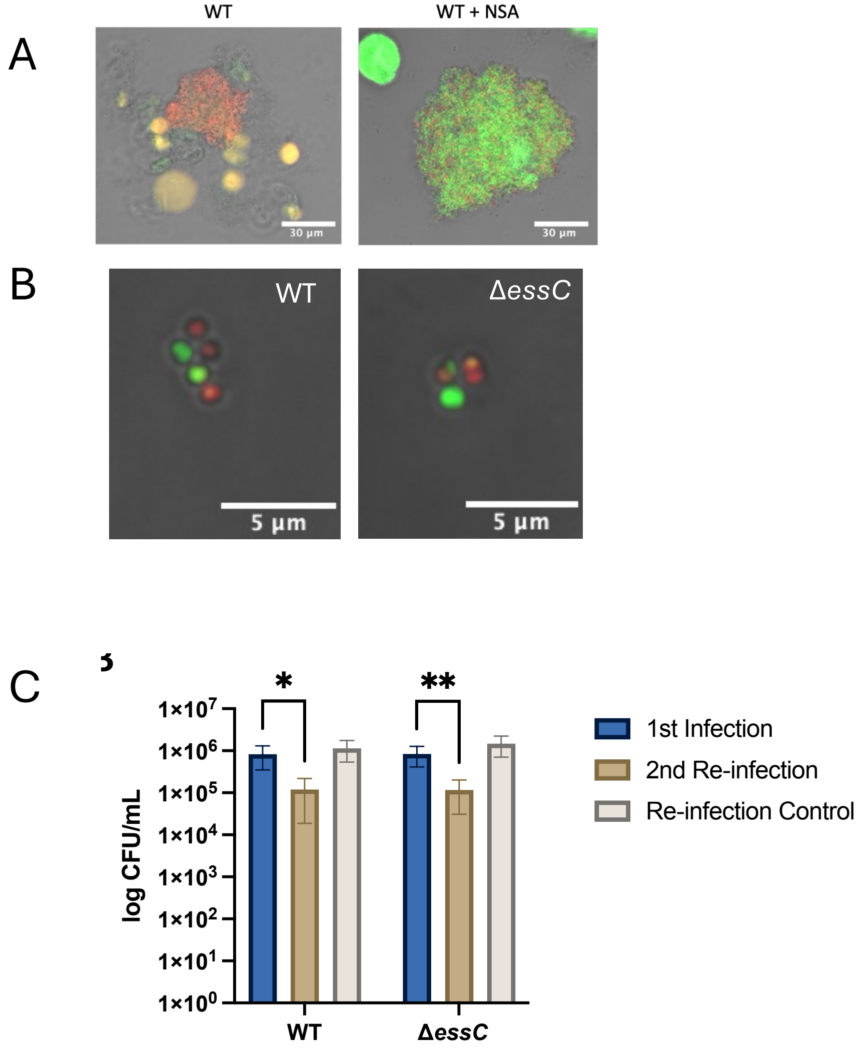


**Figure S6.** A) Representative images of microcolonies stained with LIVE/DEAD at 10 HPI of JE2 WT-infected THP-1 macrophages with and without NSA inhibitor, scale bar represents 30 µm. B) Intracellular JE2 WT collected from 6 hpi and stained with LIVE/DEAD (SYTO9 in green and propidium iodide in red) and imaged at x100 magnification, scale bar represents 5 µm. C) THP-1 macrophages infected with JE2 WT and Δ*essC* for 6 hr before lysed to collect intracellular bacteria, and lysed bacteria infected a secondary set of THP-1 macrophages for 4 hrs before lysis and CFU/ml plated. Re-infection control represents equal number of bacteria to that collected after 6 hpi but from subculture to infect secondary set of THP-1 macrophages. Mann-Whitney tests and Holm-Šídák's multiple comparison test revealed statistical significance between 1$^{st}$ and 2$^{nd}$ infection for WT and Δ*essC*, presented as (*P<0.05, **P<0.01). Data represents N=3, and graph presents ± SD.

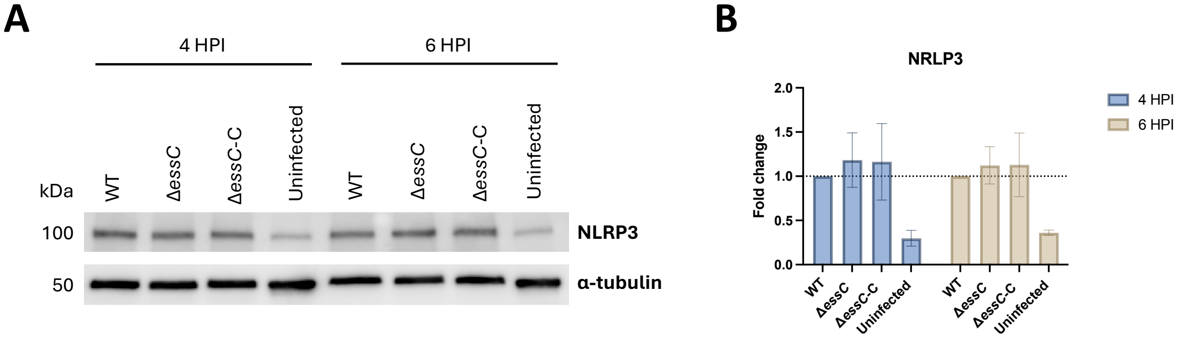


**Figure S7** BMDMs were infected with JE2 WT, Δ*essC*, and Δ*essC*-C for 1 hr at MOI 10:1 and lysates were collected at 4 and 6 hpi. A) representative immunoblots probed with anti-NLRP3 and anti-α-tubulin loading control. B) quantification of NLRP3 bands normalised to loading control, presented as fold change relative to WT at each timepoint. N=3, mean ±SD and no statistical differences between WT and mutant groups.

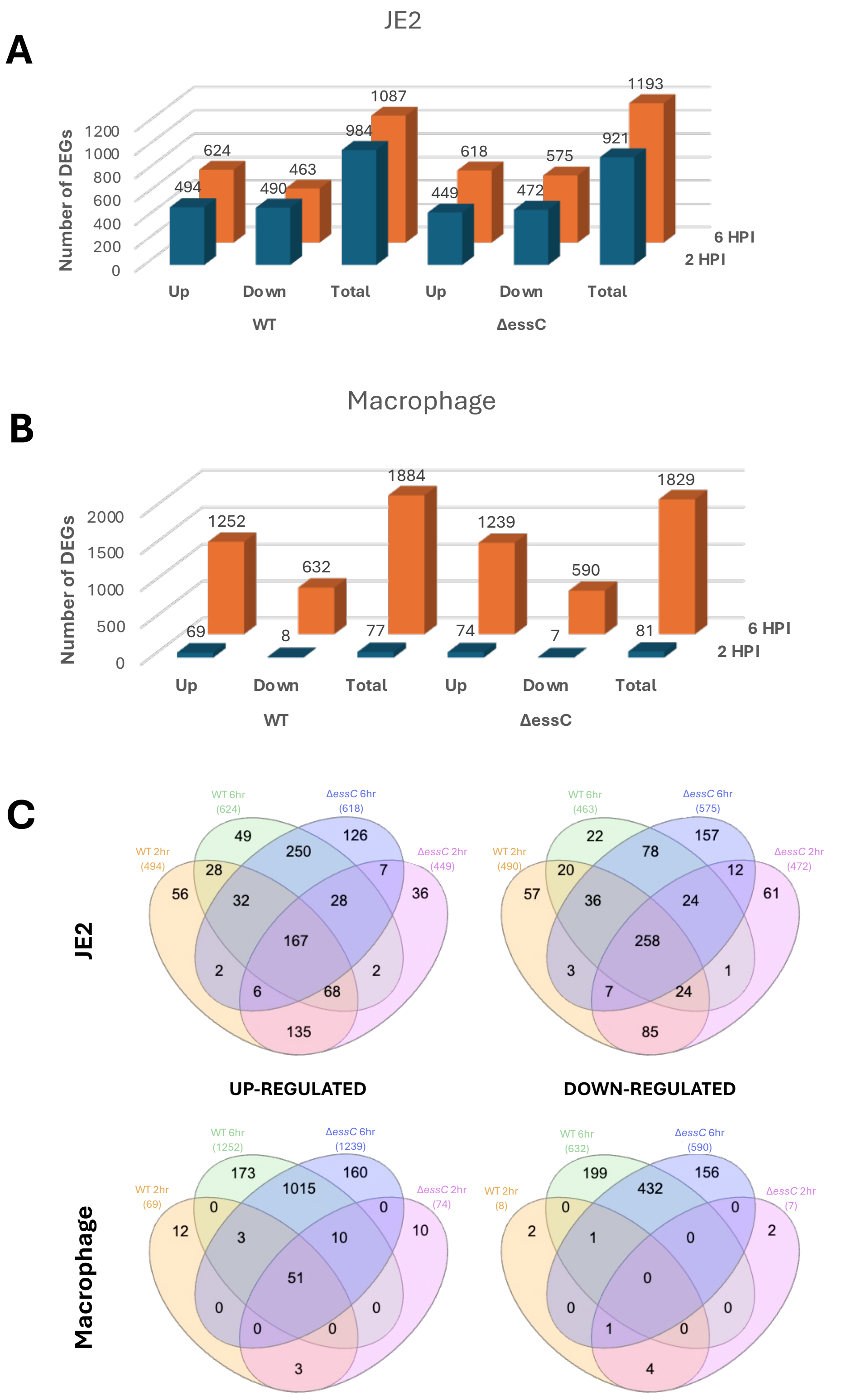


**Figure S8.** Differentially expressed genes (DEGs) were generated from DESeq2 analysis of dual RNAseq counts. **A**) The number of DEGs from comparison of JE2 *S. aureus* from infected cells to planktonic control at each timepoint for WT and Δ*essC* at 2 and 6 hpi. **B**) The number of DEGs comparing WT- or Δ*essC*-infected macrophages to uninfected macrophages at 2 and 6 hpi. DEGs were selected under the threshold of p.adj < 0.05 and log2FC ³1 and £-1. DEGs are represented at 2 hpi in blue, and at 6 hpi in orange. **C**) Leaf plots displaying the overlap of DEGs between WT and Δ*essC* condition, at 2 and 6 hpi for up- and down-regulated JE2 *S. aureus* and macrophage DEGs.

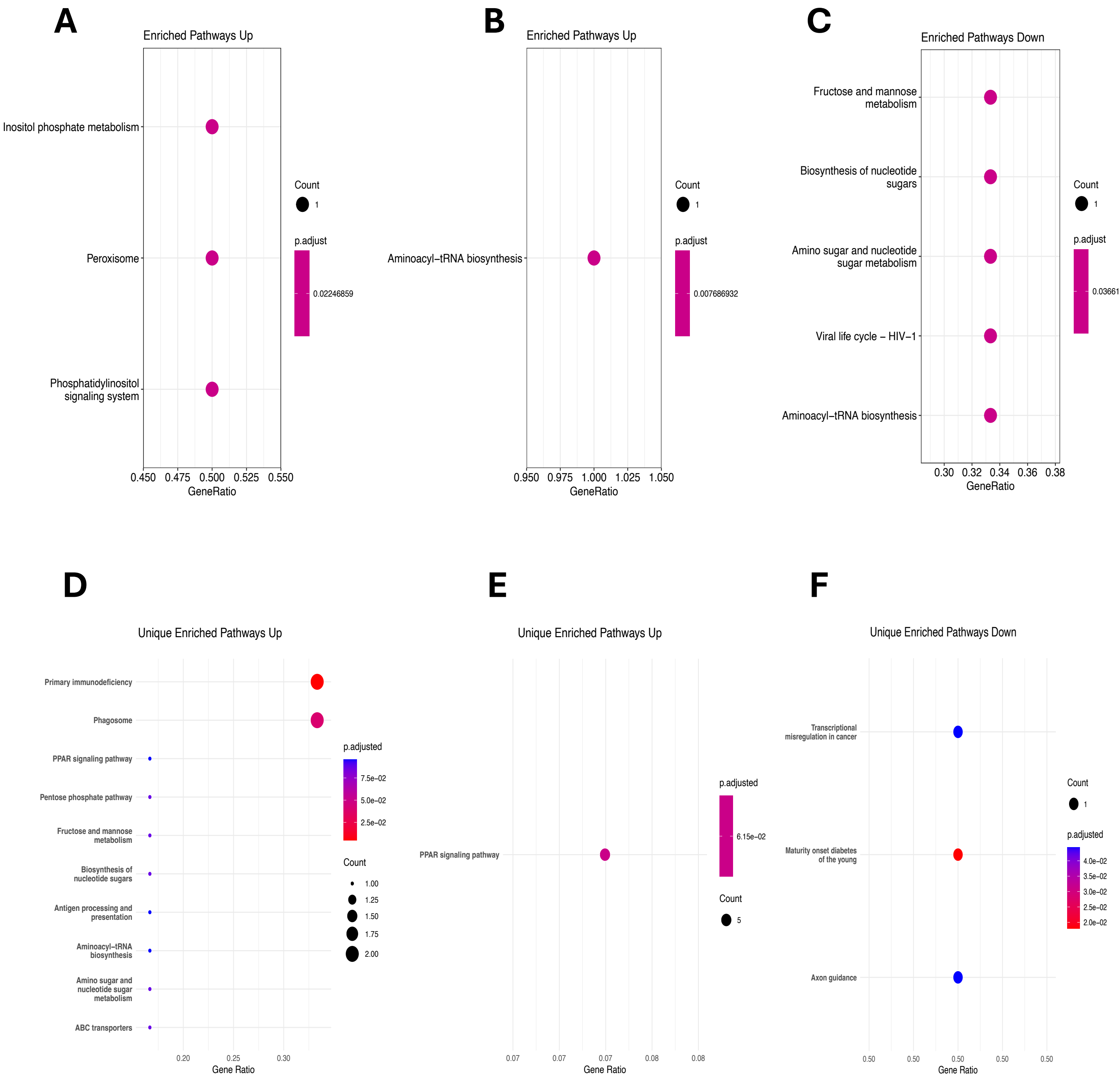


**Figure S9.** KEGG pathways enriched from Δ*essC*- vs WT-infected macrophage up- and down-regulated DEGs at 2 hpi (**A, C**) and 6 hpi (**B**). Number of DEGs identified in enriched pathway is represented by circle size (Count) and P-adjusted value (P. adjust) is represented by colour. KEGG pathway enrichment from up- and down-regulated WT-infected macrophage vs. uninfected macrophage DEGs at 2 hpi (**D, F**) and 6 hpi (**E**). Number of genes associated with the term is represented by circle size (count) and P-adjusted value is represented by colour.

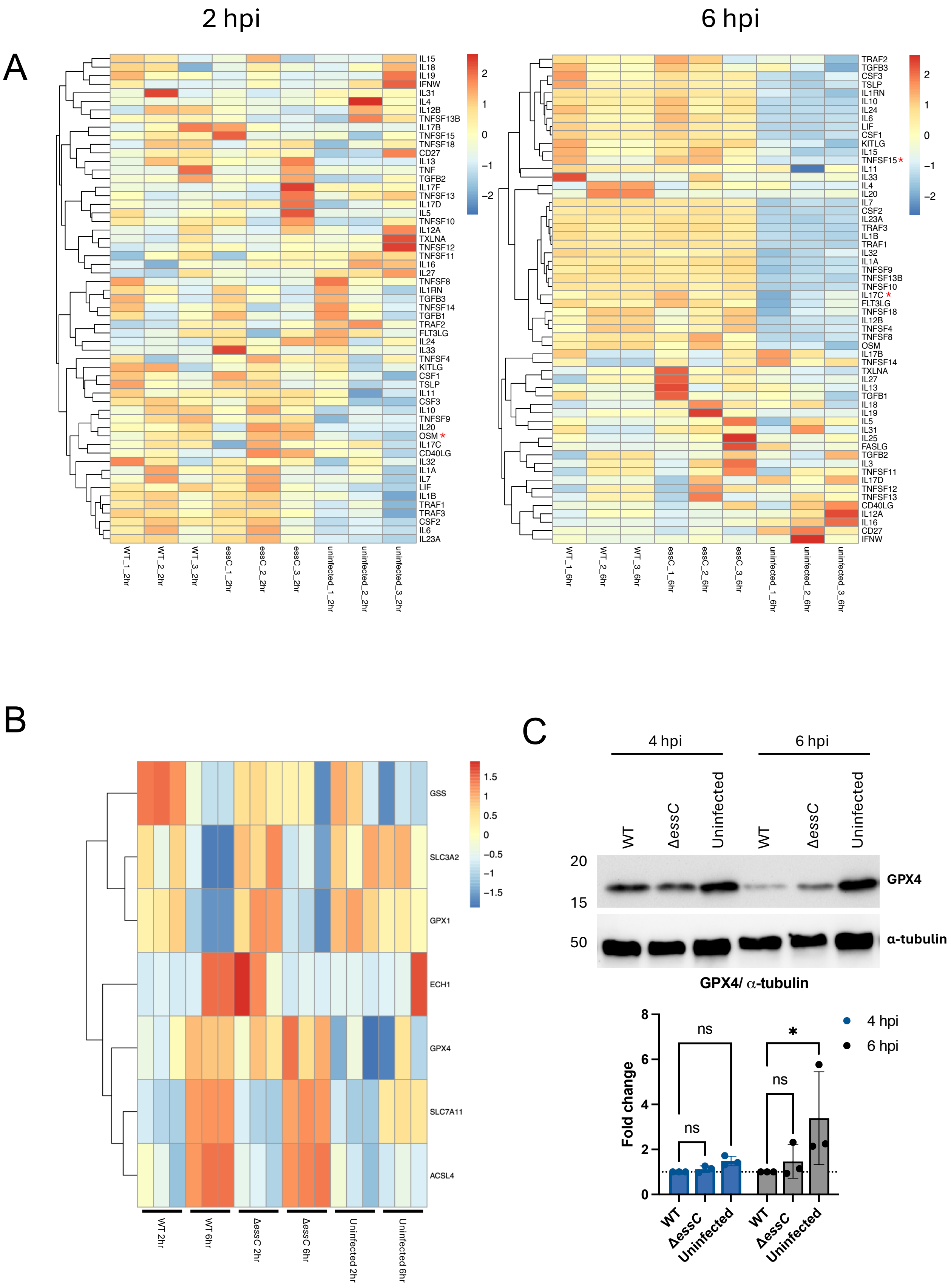


**Figure S10. A**) Heatmap of cytokine expression JE2 WT or Δ*essC* infected THP-1 macrophages and uninfected macrophages, at 2 and 6 hpi. Red asterisks indicate significantly differential gene expression between Δ*essC* and WT when compared to uninfected macrophages. **B)** Normalised macrophage counts from dual RNA-seq showing key genes in the ferroptosis pathway. **C)** THP-1 lysates obtained at 6 hpi were probed with anti-GPX4 and with anti- α-tubulin (loading control). A representative immunoblot is shown. Quantification of GPX4 bands normalised to loading control is shown below the image and presented as fold change relative to WT. N = 3, mean ± SD.

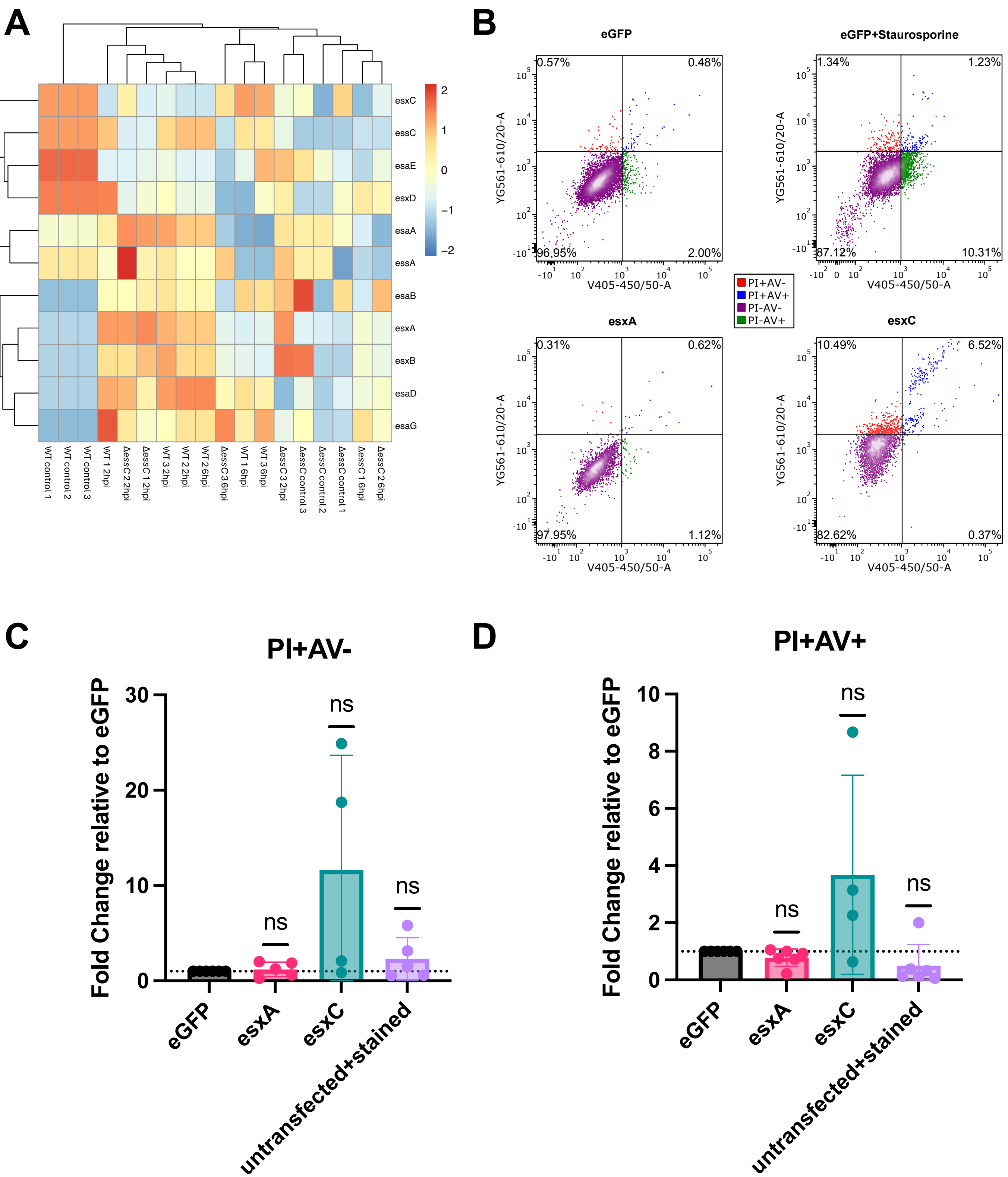


**Figure S11 A**) Heatmap of normalised expression depicting transcript abundance of T7SS locus across bacterial samples. Expression is represented as a computed Z-score denoting high (red) and low (blue) expression across samples. **B**) Representative flow cytometry plots of eGFP-positive macrophages gated for Annexin V and PI (for data shown in Figure 5D). iBMDMs were transfected with either pEGFP-N1, pEGFP-N1+*esxA*, or pEGFP-N1+*esxC* via nucleofection. At 12 hrs post transfection, macrophages were stained with Annexin V (AV) Blue Pacific and propidium iodide (PI) and analysed by flow cytometry. Transfected cells were gated for eGFP expression. The number of PI+AV- (**C**) and PI+AV+ (**D**) macrophages relative to the empty control vector (eGFP) are shown. N = 3, mean ± SD and difference were not significant (ns) by a one-sample t-test.

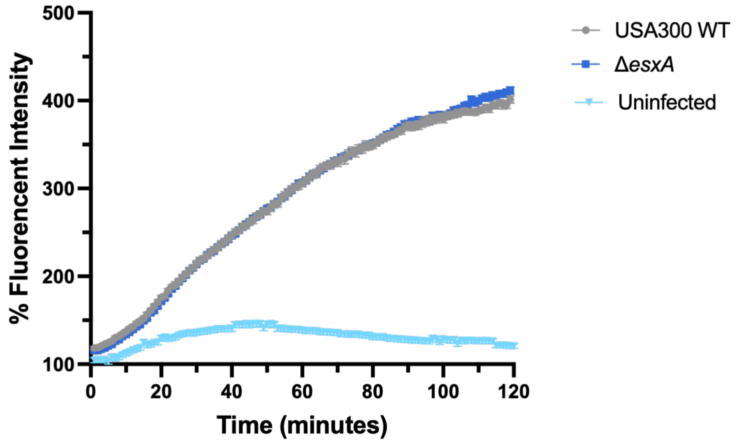


**Figure S12**. Cumulative Fluo-8 fluorescence in macrophages following infection at MOI 10:1. Macrophages were loaded with Fluo-8 AM and infected with USA300 WT or Δ*esxA.* Time-lapse fluorescence imaging was performed at 1 min intervals for 120 min. For each time point, fluorescence intensity was quantified across the entire field of view and expressed as cumulative $\% = (\Delta F/F_0) \times 100$ relative to baseline. Data represents mean ± SEM, N=3.
