## Supplementary material for "The staphylococcal type VII secretion system delays inflammatory cell death during macrophage infection": Suppl tables

**Table S1. Bacterial strains used in this study**

| Strain | Description | Source or Reference |
|---|---|---|
| *S. aureus* JE2 WT | USA300 LAC plasmid-cured | BEI Resources |
| *S. aureus* JE2 Δ*essC* | USA300 JE2 with a deletion in *essC* | Kengmo Tchoupa *et al.*, 2020 |
| *S. aureus* JE2 Δ*esxC* | USA300 JE2 with a deletion in *esxC* | Kengmo Tchoupa *et al.*, 2020 |
| *S. aureus* JE2 Δ*essC* + pOS1-*essC* (Δ*essC*-C) | Complemented JE2 Δ*essC* strain | Kengmo Tchoupa *et al.*, 2020 |
| *S. aureus* JE2 WT + pJL94-YFP | YFP-expressing JE2 WT strain | Wolz lab, University of Tubingen |
| *S. aureus* JE2 WT + pMV158-mCherry | mCherry expressing JE2 WT strain | Foster lab, University of Sheffield |
| *S. aureus* JE2 Δ*essC* + pJL94-YFP | YFP-expressing Δ*essC* strain | Wolz lab, University of Tubingen |
| *S. aureus* JE2 Δ*essC* + pMV158-mCherry | mCherry expressing Δ*essC* strain | This study |
| *S. aureus* JE2 Δ*esxC* + pMV158-mCherry | mCherry expressing Δ*esxC* strain | This study |
| *S. aureus* JE2 WT + pMV158-GFP | GFP-expressing JE2 WT strain | Foster lab, University of Sheffield |
| *S. aureus* JE2 Δ*essC* + pMV158-GFP | GFP-expressing Δ*essC* strain | This study |
| *S. aureus* USA300 WT | USA300 LAC strain | Schneewind lab, University of Chicago |
| *S. aureus* USA300 Δ*esxA* | USA300 LAC with a deletion in *esxA* | Kengmo Tchoupa *et al.*, 2020 |
| *S. aureus* USA300 WT + pMV158-mCherry | mCherry expressing USA300 WT strain | This study |
| *S. aureus* USA300 Δ*esxA* + pMV158-mCherry | mCherry expressing Δ*esxA* strain | This study |
| *E. coli* DH5α | NEB 5-alpha Competent *E. coli* | NEB |
| *E. coli* DC10B | DNA cytosine methyltransferase mutant | Monk et al., 2012, Foster lab, Trinity College Dublin |
| *E. coli* DH5α + pEGFP-N1 | Expressing mammalian plasmid for eGFP | Royle lab, University of Warwick |

**Table S2. Primers used in this study**

| Primer Name | Primer Sequence (5'-3') | Description |
|---|---|---|
| EssC_comp_RBS_Pst1_FP | GCGCTGCAGTTGAGAGGAGAGAAAATGCATAAATTGATTATAAA | Insertion of *essC* into pOS1 for Δ*essC* complement |
| EssC_comp_Sma1_RP | GCGGCGCCCGGGCTATTTAAACCATCTAATCT | Insertion of *essC* into pOS1 for Δ*essC* complement |
| CMV_Nhe1_esxC_FP | GCGGCTAGCATGAATTTTAATGATATTGA | Insertion of *esxC* into CMV containing eGFP |
| esxC_Age1_RP | ACTACCGGTGTATTCATTGCTTTATTAAAAT | Insertion of *esxC* into CMV containing eGFP |

### Table S3 Δ*essC*-infected vs. WT-infected macrophage DEGs

| | Up-regulated | | | | Down-regulated | | | |
|---|---|---|---|---|---|---|---|---|
| Time-point | Gene ID | Gene Description | log2FC | P-adjusted | Gene ID | Gene Description | log2FC | P-adjusted |
| 2 hpi | ECH1 | Enoyl-CoA Hydratase 1 | 4.63675273 | 0.00631309 | VARS1 | valyl-tRNA synthetase 1 | -7.3773693 | 0.00016684 |
| | ITPK1 | inositol-tetrakisphosphate 1-kinase | 3.45003766 | 0.0200537 | FLII | FLII actin remodelling protein | -4.0271268 | 2.63E-06 |
| | LENG8 | Leukocyte receptor cluster member 8 | 3.25622419 | 0.03661222 | SMARCB1 | SWI/SNF related, matrix associated, actin dependent regulator of chromatin, subfamily b, member 1 | -3.7073857 | 0.02626993 |
| | GARIN4 | golgi associated RAB2 interactor family member 4 | 1.07479078 | 0.00578814 | GFUS | GDP-L-fucose synthase | -3.7058334 | 2.63E-06 |
| 6 hpi | LENG8 | leukocyte receptor cluster member 8 | 6.16262265 | 0.00380677 | YWHAE | tyrosine 3-monooxygenase/tryptophan 5-monooxygenase activation protein epsilon | -6.1386573 | 0.01265654 |
| | VARS1 | valyl-tRNA synthetase 1 | 6.13225563 | 0.00773871 | HNRNPL | heterogeneous nuclear ribonucleoprotein L | -6.0057683 | 0.00027893 |
| | DENND11 | DENN domain containing 11 | 6.04084603 | 0.03258711 | AL593848.2 | Uncharacterised | -5.7549737 | 0.01900565 |
| | RRN3 | RRN3 homolog, RNA polymerase I transcription | 3.64188183 | 0.03258711 | ADCK5 | aarF domain containing kinase 5 | -5.5110948 | 0.00051436 |
| | | | | | CUTA | cutA divalent cation tolerance homolog | -5.2794981 | 2.71E-06 |
| | | | | | ENSG00000292221 | Uncharacterised | -5.0503036 | 0.0007242 |
| | | | | | RPL7A | ribosomal protein L7a | -3.8553438 | 0.04846518 |
| | | | | | UQCRC2 | ubiquinol-cytochrome c reductase core protein 2 | -3.8271977 | 0.0262937 |
| | | | | | TDO2 | tryptophan 2,3-dioxygenase | -3.6545241 | 0.01112199 |
| | | | | | KANSL1 | KAT8 regulatory NSL complex subunit 1 | -3.6382075 | 0.00770905 |
| | | | | | PRRC2A | proline rich coiled-coil 2A | -3.5478705 | 0.00690971 |
| | | | | | PPFIA1 | PTPRF interacting protein alpha 1 | -3.2848476 | 0.00770905 |
| | | | | | LSM14A | LSM14A mRNA processing body assembly factor | -3.2132807 | 0.00108275 |
| | | | | | CC2D1A | coiled-coil and C2 domain containing 1A | -3.1846834 | 2.71E-06 |
| | | | | | H2AC19 | H2A clustered histone 19 | -1.6917884 | 0.03258711 |

### Table S5. Δ*essC* vs. WT DEGs during intracellular infection

| Up-regulated | Down-regulated |
|---|---|

| Time-point | Gene ID | Gene Description | log2FC | P-adjusted | Gene ID | Gene Description | log2FC | P-adjusted |
|---|---|---|---|---|---|---|---|---|
| 2 hpi | B7H15_04460 | cold-shock protein; CspC | 1.59488747 | 3.24E-09 | B7H15_01565 | essC | -6.800493496 | 1.34E-05 |
| | B7H15_15050 | cold-shock protein; CspB | 1.439942864 | 0.000378718 | B7H15_01030 | ornithine aminotransferase 1 (OatA) | -1.114292957 | 0.04842354 |
| 6 hpi | B7H15_14650 | clumping factor B | 1.978519519 | 1.71E-07 | B7H15_01565 | essC | -8.196429737 | 6.81E-08 |
| | B7H15_15050 | cold-shock protein; CspB | 1.689314381 | 0.029311791 | B7H15_11145 | transcriptional regulator | -1.538409094 | 0.02931179 |
| | B7H15_14275 | CHAP domain-containing protein; SsaA | 1.647660772 | 0.020617218 | | | | |
| | B7H15_02375 | hypothetical protein; DabA | 1.087883203 | 0.009190109 | | | | |

**Table S6. Bacterial iron and oxidative stress related DEGs during macrophage infection**

Red cells highlight significantly up-regulation where P.adjusted <0.05 & Log2FC > 1 and blue cells highlight down-regulation where P.adjusted <0.05 & Log2FC -1 <

| | | 2 hpi | | | | 6 hpi | | | | |
|---|---|---|---|---|---|---|---|---|---|---|
| | | WT | | *ΔessC* | | WT | | *ΔessC* | | |
| GENE ID | GENE DESCRIPTION | log2Fc | p.adjusted | log2Fc | p.adjusted | log2Fc | p.adjusted | log2Fc | p.adjusted | KEY ROLE |
| **B7H15_07835** | Ferredoxin | 6.28020549 | 0.00020012 | 1.15218333 | 0.45734825 | 7.02342305 | 4.19E-07 | 8.25357282 | 2.42E-05 | Iron metabolism |
| **B7H15_12675** | ferrichrome ABC transporter substrate-binding protein | 2.15476146 | 2.12E-05 | 0.93039126 | 0.20541051 | 2.57610288 | 6.48E-10 | 3.29292611 | 9.99E-17 | Iron metabolism |
| **B7H15_05885** | hemin ABC transporter permease | 1.70265397 | 5.77E-09 | 0.80588805 | 0.01658917 | 2.96788113 | 1.82E-15 | 2.41813261 | 3.40E-11 | Iron import |
| **B7H15_10380** | peroxiredoxin | 1.30527014 | 0.00057479 | 0.71091911 | 0.18075335 | 0.34454752 | 0.61400646 | -0.1015631 | 0.90029615 | Antioxidant |
| **B7H15_06800** | glutathione peroxidase | 1.22462092 | 0.02898216 | 1.03456736 | 0.05979316 | 0.60059176 | 0.16180451 | -0.0392415 | 0.93674638 | Antioxidant |
| **B7H15_00640** | iron-siderophore ABC transporter permease | 1.14678226 | 1.80E-08 | 0.71515295 | 0.00179877 | 3.39387468 | 7.64E-10 | 3.77893354 | 4.38E-12 | Iron import |
| **B7H15_12175** | NADP-dependent oxidoreductase | 1.12609223 | 1.14E-07 | 0.87655862 | 6.27E-05 | 0.60504865 | 0.20783963 | 0.40113744 | 0.90188383 | Antioxidant |
| **B7H15_00935** | monooxygenase IsdI | 1.05004006 | 0.33892074 | 2.56305938 | 0.00106381 | 1.14345836 | 0.15278937 | 0.96789882 | 0.2200492 | Iron metabolism |
| **B7H15_07515** | peptide-methionine (S)-S-oxide reductase | 2.28539504 | 4.65E-34 | 1.76695802 | 2.16E-17 | 1.44665891 | 0.01784443 | 0.29158948 | 0.68291961 | Antioxidant |
| **B7H15_14105** | thiol reductase thioredoxin | 2.72003992 | 8.06E-22 | 2.63660045 | 4.19E-18 | 1.4382813 | 0.0027836 | 0.9666057 | 0.05328204 | Antioxidant |
| **B7H15_13015** | 3-hydroxyacyl-CoA dehydrogenase | 1.49864019 | 6.49E-09 | 1.65164235 | 2.14E-11 | 1.40769303 | 9.19E-06 | 0.83578201 | 0.01034536 | Fatty acid metabolism / oxidative stress |
| **B7H15_07175** | peptide-methionine (S)-S-oxide reductase | 1.91727165 | 4.64E-11 | 1.62349113 | 5.05E-07 | 1.10394625 | 0.01592534 | 0.1786165 | 0.75189163 | Antioxidant |
| **B7H15_05625** | NrdH-redoxin | 2.20517434 | 0.00022099 | 1.15045372 | 0.08824613 | 2.12749543 | 0.00081203 | 1.36841678 | 0.03362763 | Antioxidant |
| **B7H15_13145** | oxidoreductase | 0.16107311 | 0.68285125 | 0.41989448 | 0.20598226 | 1.09563699 | 0.00658524 | 0.99511173 | 0.01254681 | Antioxidant |
| **B7H15_06725** | 2-oxoglutarate ferredoxin oxidoreductase subunit alpha | 1.67377195 | 8.62E-10 | 1.26813865 | 1.22E-05 | 1.07591461 | 0.00010407 | 0.92029698 | 0.00064851 | Iron metabolism |
| **B7H15_04745** | nitronate monooxygenase | 1.52334726 | 0.17130537 | -1.6115151 | 0.3107784 | 2.13486489 | 0.00165534 | 0.33101247 | 0.65755944 | Oxidative stress |
| **B7H15_14185** | FeoB-associated Cys-rich membrane protein | -4.51611633 | 0.00110032 | -2.6906529 | 0.06853076 | -2.9970937 | 0.00031598 | -3.6768134 | 0.00768548 | Iron import |
| **B7H15_07045** | LexA repressor | -1.48806839 | 5.34E-09 | -0.9276486 | 0.00032039 | -2.1234788 | 5.09E-16 | -2.0107253 | 3.62E-15 | Stress |
| **B7H15_00470** | oxidoreductase | -1.40873348 | 0.00013061 | -0.5819172 | 0.14112319 | -1.9458927 | 4.47E-05 | -2.807869 | 1.79E-08 | Antioxidant |
| **B7H15_01285** | nitric oxide dioxygenase | -1.36957397 | 2.91E-09 | -0.4978926 | 0.02885503 | -1.1324639 | 1.43E-11 | -0.6060304 | 0.00018553 | Stress |
| **B7H15_14190** | ferrous iron transporter B | -2.46767475 | 0.00141233 | -1.8207808 | 0.04148644 | -1.3518167 | 0.0094025 | -0.228685 | 0.69467668 | Iron import |
| **B7H15_05480** | quinol oxidase subunit 3 | -1.52908369 | 1.75E-13 | -0.9015857 | 2.62E-05 | -1.341004 | 1.35E-08 | -0.6588341 | 0.00705129 | Oxidative stress |
| **B7H15_01775** | NADH-dependent flavin oxidoreductase | -2.4877857 | 6.12E-17 | -2.8161739 | 2.17E-16 | -1.090594 | 0.00431551 | -0.7132488 | 0.06459346 | ROS regulation |
| **B7H15_05475** | quinol oxidase subunit 4 | -1.3790727 | 1.35E-08 | -0.8744508 | 0.00055889 | -1.048649 | 0.00017087 | -0.4270348 | 0.15001266 | Oxidative stress |
| **B7H15_03845** | oxidoreductase | -0.67507298 | 0.00058956 | -0.0655682 | 0.78726952 | -1.0185939 | 1.08E-06 | -0.4307523 | 0.04248121 | Antioxidant |
| **B7H15_00035** | NAD(P)H-hydrate dehydratase | -1.11882987 | 4.76E-05 | -0.5606767 | 0.04618542 | 0.28571766 | 0.55184364 | 1.51345238 | 0.0001243 | ROS regulation |
| **B7H15_12570** | MFS transporter | -1.48238782 | 0.02947655 | -0.1169876 | 0.84304628 | 1.39623613 | 0.06490696 | 1.50876954 | 0.0391018 | Iron import |
| **B7H15_04715** | iron-sulfur cluster assembly scaffold protein NifU | 0.14910506 | 0.91408613 | 0.17789425 | 0.88282603 | 0.78198018 | 0.33539639 | 1.9255865 | 0.00452921 | Iron metabolism |
| **B7H15_02720** | molecular chaperone Hsp33 | 0.50818083 | 0.50573882 | 0.10630545 | 0.9043957 | 0.96891508 | 0.06232972 | 1.47342483 | 0.00121686 | Oxidative stress |
| **B7H15_06330** | peptide deformylase | -0.64628009 | 0.58023553 | 0.38686927 | 0.72097795 | 0.14266063 | 0.83500074 | 1.41181695 | 0.0057955 | Iron metabolism |
| **B7H15_12110** | iron-siderophore ABC transporter permease | 0.40995115 | 0.15278357 | 0.34185646 | 0.26090491 | 0.93183182 | 0.00045781 | 1.37089938 | 5.12E-08 | Iron import |
| **B7H15_12105** | iron-dicitrate transporter subunit FecD | -0.05778805 | 0.87961612 | 0.09857789 | 0.76040991 | 0.55473379 | 0.10762875 | 1.20928333 | 0.00013304 | Iron import |
| **B7H15_06680** | 3-oxoacyl-ACP reductase | -0.80973102 | 0.02489242 | -1.1591539 | 0.00266757 | 0.74968851 | 0.03468837 | 1.18444619 | 0.00034735 | Fatty acid metabolism / oxidative stress |
| **B7H15_08540** | metal ABC transporter permease | -0.36080893 | 0.16585417 | 0.118282 | 0.6699897 | 0.46038749 | 0.20289897 | 1.15783584 | 0.00028475 | Metal ion import |
| **B7H15_07015** | catalase | 1.32509921 | 4.50E-17 | 0.9593473 | 3.81E-09 | 0.11506959 | 0.91098258 | -1.6314732 | 0.0489839 | Antioxidant |
| **B7H15_11120** | oxidoreductase | -0.97502265 | 0.00013024 | -1.233838 | 4.55E-06 | -0.9891682 | 1.25E-07 | -0.5506947 | 0.00124991 | Antioxidant |
| **B7H15_11900** | oxidoreductase | 0.13419389 | 0.87193129 | -1.0272259 | 0.23489408 | -0.7064619 | 0.31446677 | -1.6072105 | 0.02046925 | Antioxidant |
| **B7H15_04565** | thiol reductase thioredoxin | -0.16942967 | 0.541848 | -0.1865176 | 0.51583976 | -0.812051 | 0.03062084 | -1.2379547 | 0.00062091 | Antioxidant |
| **B7H15_04840** | FAD-dependent oxidoreductase | 0.51543052 | 0.00575493 | 0.55707789 | 0.00285521 | -0.3209207 | 0.59209606 | -1.1452235 | 0.02657201 | Antioxidant |
| **B7H15_04705** | Fe-S cluster assembly protein SufD | -0.63358908 | 0.0023946 | -0.8800024 | 1.79E-05 | -0.7787702 | 0.0130101 | -1.1405314 | 0.00014761 | Iron import |
| **B7H15_06190** | laccase | 0.06418214 | 0.75435248 | -0.0866664 | 0.65717131 | -0.3440057 | 0.22995313 | -1.0043074 | 0.00010535 | Oxidative stress |

### Table S7. Infected macrophage ferroptosis DEGs

| | 2 hpi | | | | 6 hpi | | | |
|---|---|---|---|---|---|---|---|---|
| | WT | | *ΔessC* | | WT | | *ΔessC* | |
| Gene | log2FC | p.adj | log2FC | p.adj | log2FC | p.adj | log2FC | p.adj |
| **SLC7A11** | 0.19328903 | 0.94067202 | 0.18910356 | 0.99997432 | 1.09197029 | 1.61E-23 | 1.04183767 | 2.63E-21 |
| **SLC3A2** | -0.0474056 | 0.95561169 | 0.06541643 | 0.99997432 | -0.3554818 | 0.00150079 | -0.2724769 | 0.02093726 |

| | | | | | | | | |
|---|---|---|---|---|---|---|---|---|
| **GPX4** | 0.12518472 | 0.7565388 | 0.21473499 | 0.34860419 | 0.20969721 | 0.07764647 | 0.23102059 | 0.04929803 |
| **ACSL4** | 0.08512201 | 0.96098498 | 0.01667168 | 0.99997432 | 1.02044145 | 5.68E-29 | 0.98898304 | 5.84E-27 |
| **GSS** | 0.13552874 | 0.7552555 | 0.02088449 | 0.99997432 | 0.06408781 | 0.72305187 | 0.09765643 | 0.56839589 |
| **GPX1** | -0.1473183 | 0.65103729 | 0.0398521 | 0.99997432 | -0.6112209 | 6.40E-08 | -0.5010597 | 1.50E-05 |
| **ECH1** | 0.79884405 | 0.95253959 | 5.43559678 | 0.00259521 | 0.62944161 | 0.85657949 | -4.6911728 | 0.09399587 |
