## Supplementary material for "The staphylococcal type VII secretion system delays inflammatory cell death during macrophage infection": Suppl methods

**Supplementary Methods**

**Bacterial strains and culture conditions:** *S. aureus* strains were grown in tryptic soya broth (TSB) (Merck) aerobically. Complemented strains of *S. aureus* were grown in TSB supplemented with 10 µg/ml chloramphenicol. Δ*essC* and Δ*esxC* were generated via allelic replacement from previous studies[1,2]. Complementation of Δ*essC* was achieved via amplification of *essC* from USA300 JE2 genomic DNA and inserted into pOS1 by restriction digestion. pOS1*essC* was transformed into *E. coli* DC10B by heat shock, and subsequently by electroporation into intermediate *S. aureus* strain RN4220 and then electroporated into JE2 strains.

**Macrophage culture:** Human monocyte THP-1s (ATCC) were cultured in suspension at 37°C 5% $CO_2$ in Roswell Park Memorial Institute (RPMI) media (Gibco) supplemented with 10% fetal bovine serum (FBS) (Gibco), 4.5 g/L D-glucose, 2.383 g/L HEPES buffer, L-glutamine, 1.5 g/L sodium bicarbonate, 110 mg/L sodium pyruvate (ATCC modification). THP-1 monocytes were differentiated with 50 ng/ml phorbol-12-myrisate-13-acetate (PMA) (Sigma) in RPMI for 72 hrs followed by an overnight rest without PMA. Bone marrow derived macrophages (BMDMs) were extracted from 8-week old C57BL/6 femur tibia and cultured for 7 days in RPMI (ATCC modification) + 10% FBS + 3% PenStrep containing 0.25 ng/ml M-CSF (BioLegend) and used in experiments on day 8. iBMDMs were kindly gifted from Dr Avinash Shenoy (Imperial College London) and were maintained in RPMI ATCC modification + 0.25 ng/ml M-CSF until seeding for experiments where M-CSF was removed.

**Measuring intracellular bacterial damage and reinfection assays**: Intracellular *S. aureus* infection of THP-1 macrophages was carried out with refreshment in lysostaphin after extracellular killing as described above. At 6 hpi, THP-1 macrophages were lysed to obtain intracellular bacteria by the addition of cold H20 and CFU/ml obtained. Intracellular bacteria collected at 6 hpi were then resuspended in RPMI +10% FBS and used to infect a new set of THP-1 macrophages as previously described. At 4 hr post- “secondary”-infection, THP-1 macrophages were lysed with cold water and intracellular CFU/ml obtained.

Intracellular *S. aureus* obtained from the first 6 hr infection were stained with LIVE/DEAD BacLight Bacterial Viability Kit (ThermoFisher Scientific, United States) according to manufacturer's instructions and imaged on Leica BMi8 inverted widefield fluorescent microscope with ORCA-Flash4.0 V2 digital CMOS camera (Hamamatsu Photonics, Japan).

**Dual RNA-seq analysis**

Gene ontology analysis was carried out by first generating 'GOterms' mapped to the JE2 genome using InterProScan. Then, GOenrichment was performed using the clusterProfiler 'R' package (version 4.10.1) and 'enrichGO' function[3,4]. KEGG enrichment analysis was also carried out using the 'enrichKEGG' function from the clusterProfiler package[5]. Both enrichGO and enrichKEGG were carried out with a p-value cutoff of 0.05 and the p.adjusted method set to Benjamini-Hochberg (BH).
